# DDX5 (p68) orchestrates β-catenin, RelA and SP1 mediated MGMT gene expression: Implication in TMZ chemoresistance

**DOI:** 10.1101/2023.04.29.538802

**Authors:** Rajni Shaw, Subhajit Karmakar, Malini Basu, Mrinal K. Ghosh

## Abstract

DDX5 (p68) upregulation has been linked with various cancers of different origins, especially Colon Adenocarcinomas. Similarly, across cancers, MGMT has been identified as the major contributor of chemoresistance against DNA alkylating agents like Temozolomide (TMZ). TMZ is an emerging potent chemotherapeutic agent across cancers under the arena of drug repurposing. Recent studies have established that patients with open MGMT promoters are prone to be innately resistant or acquire resistance against TMZ compared to its closed conformation. However, not much is known about the transcriptional regulation of MGMT gene in the context of colon cancer. This necessitates studying MGMT gene regulation, which directly impacts the cellular potential to develop chemoresistance against alkylating agents. Our study aims to uncover an unidentified mechanism of DDX5-mediated MGMT gene regulation. Experimentally, we found that both mRNA and protein expression levels of MGMT were elevated in response to p68 overexpression in multiple human colon cancer cell lines and vice-versa. Since p68 cannot directly interact with the MGMT promoter, transcription factors *viz*., β-catenin, RelA (p65) and SP1 were also studied as reported contributors. Through co-immunoprecipitation and GST-pull-down studies, p68 was established as an interacting partner of SP1 in addition to β-catenin and NF-κB (p50-p65). Mechanistically, luciferase reporter and chromatin-immunoprecipitation assays demonstrated that p68 interacts with the MGMT promoter *via* TCF4-LEF, RELA and SP1 sites to enhance its transcription. To the best of our knowledge, this is the first report of p68 as a transcriptional co-activator of MGMT promoter and our study identifies p68 as a novel and master regulator of MGMT gene expression.

## Introduction

In the modern era, cancer treatment is a question in itself due to variable responses by different patients to each cancer therapy. The most widely used basic form of cancer treatment is chemotherapy. Chemotherapy aggressively targets the entire body, thus expected to combat cancer malignancy and it also prepares the patient for more specific treatments. But, apart from the side effects of chemotherapy cancer patients have to suffer with chemoresistance, when the body decides to respond or not respond to chemotherapy. Defeating intrinsic and acquired drug resistance is a major obstacle in treating cancer patients because chemoresistance causes recurrence, cancer dissemination and death.

DNA repair system has several mechanisms that remove damaged portions of DNA. Cancer cells exploit these repair mechanisms to remove replication blocks created by chemotherapy. One such example of a DNA repair protein is MGMT (O-6-Methylguanine-DNA methyltransferase). MGMT: a suicide enzyme, alias alkyltransferase, represents a one-step repair mechanism, which is responsible for the removal of alkyl groups from the O6-position of guanine and the O4-position of thymine.^1^ It was found that MGMT was overexpressed for the development of cancer chemoresistance *via* Wnt,^2^ NF-κB,^3^ and SP1^4, 5^ pathways, whose inhibition downregulated MGMT expression and restored the chemosensitivity to DNA-alkylating drugs like Temozolomide. The question of induction of MGMT is highly important because of its high impact on cancer therapy, as tumors like glioblastoma and metastatic melanoma are treated with O6-alkylating agents (temozolomide, dacarbazine, chloroethylating nitrosoureas) against which MGMT offers defence. In recent years, MGMT has been identified as a major culprit of Temozolomide (TMZ) mediated chemoresistance in different cancers (e.g., colon cancer, breast cancer, glioma, etc.).^6^

Many oncogenic pathways and specific molecules have been identified in recent years that act as regulators of DNA repair genes and hence chemoresistance. One such molecule with tremendous oncogenic potential is DDX5. DDX5 or p68 is a DEAD-box protein, characterized by the conserved motif Asp-Glu-Ala-Asp (DEAD) and identified as a putative RNA helicase.^7, 8^ P68 expression is upregulated in colorectal, breast, and prostate cancers and has been identified to correlate with cell proliferation, tumor progression and transformation.^9–12^ Recent reports have established p68 interaction with transcription-related factors such as p300, RNA polymerase II, histone deacetylase, Smad3, cAMP-response element-binding protein, and others.^11, 13–16^ It also acts as a commanding co-activator of tumor suppressor p53 as well as NFκB (p50-p65), MyoD, and β-catenin.^10–15, 17–19^ Moreover, studies have shown p68 recruitment to the promoters of responsive genes under conditions in which these transcription factors are activated,^10, 12, 17^ consistent with a role in transcription initiation. P68 does not have a DNA-binding domain. Hence, it acts as a strong co-activator for many important signalling molecules and amplifies their effect by multiple folds. Thus, it would be intriguing to explore the novel involvement of DDX5 (p68) as a co-activator in the transcriptional activation of MGMT in the arena of TMZ-mediated chemoresistance in colon cancer. It would be interesting to investigate the molecular mechanisms defining DDX5 (p68) as a master regulator of MGMT, and thus find out possible therapeutic approaches for cancer chemotherapy to combat chemoresistance.

## Results

### Concomitant gain of DDX5 (p68) along with β-catenin, RelA, SP1 and MGMT expressions in human colon cancer

To understand the connection between MGMT and DDX5 (p68) in colon cancer, we conducted immunohistochemical analysis in colon carcinoma (n = 20) and adjacent normal tissue samples (n = 05). MGMT and p68 showed sharp elevated expressions in colon carcinoma tissues as compared to the normal samples. As expected, β-catenin, RelA and SP1 also show enhanced expressions in carcinoma samples (Fig. 1A). Upon quantification by H-scoring, p68, MGMT, β-catenin, RelA and SP1 depicted significant differences in staining intensities between colon carcinoma and normal samples (Fig. 1B & 1C). The variation in staining intensities (measured by H-scores) of p68, MGMT, β-catenin, RelA and SP1 between normal and colon carcinoma samples (represented by Box Plot) were statistically significant and was predicted by the Mann–Whitney U test analysis (Fig. 1D). Moreover, the dissimilarity between the average H-scores of MGMT, p68, β-catenin, RelA and SP1 in colon carcinoma samples was statistically significant compared to the normal samples (Fig. 1E). H-scores of MGMT and p68 were found to bear strong positive correlation (rs = 0.8046), when statistically analysed by Spearman’s rank correlation test (Fig. 1F) as compared to the correlations between MGMT and β-catenin (rs = 0.5004); MGMT and RelA (rs = 0.6675) and MGMT and SP1 (rs = 0.7110). Additionally, we monitored the endogenous expressions of p68, MGMT, β-catenin, RelA and SP1 in HEK293 and colon cancer cell lines. We observed prominent expressions of all these molecules in all four human colon cancer cell lines (Fig. 1G) and three of them *viz.,* HCT116, SW480 and HT29 were used in further studies. Next, by using GeneMANIA platform in Cytoscape we analysed the association of DDX5 (p68), MGMT, CTNNB1 (β-catenin), RELA (p65) and SP1 along with other genes, based on their co-localization, co-expression profiles and genetic interactions (Fig. 1H). Therefore, with deep interest, we further assessed mRNA expressions of DDX5, MGMT, CTNNB1, RELA and SP1 in primary tumor vs normal tissue sample using colon adenocarcinoma (COAD) data-set, and represented as box plots [Supplementary figure S1]. Altogether, our results clearly indicate conceivable interconnections amongst p68, β-catenin, RelA, SP1 and MGMT.

**Fig. 1.**
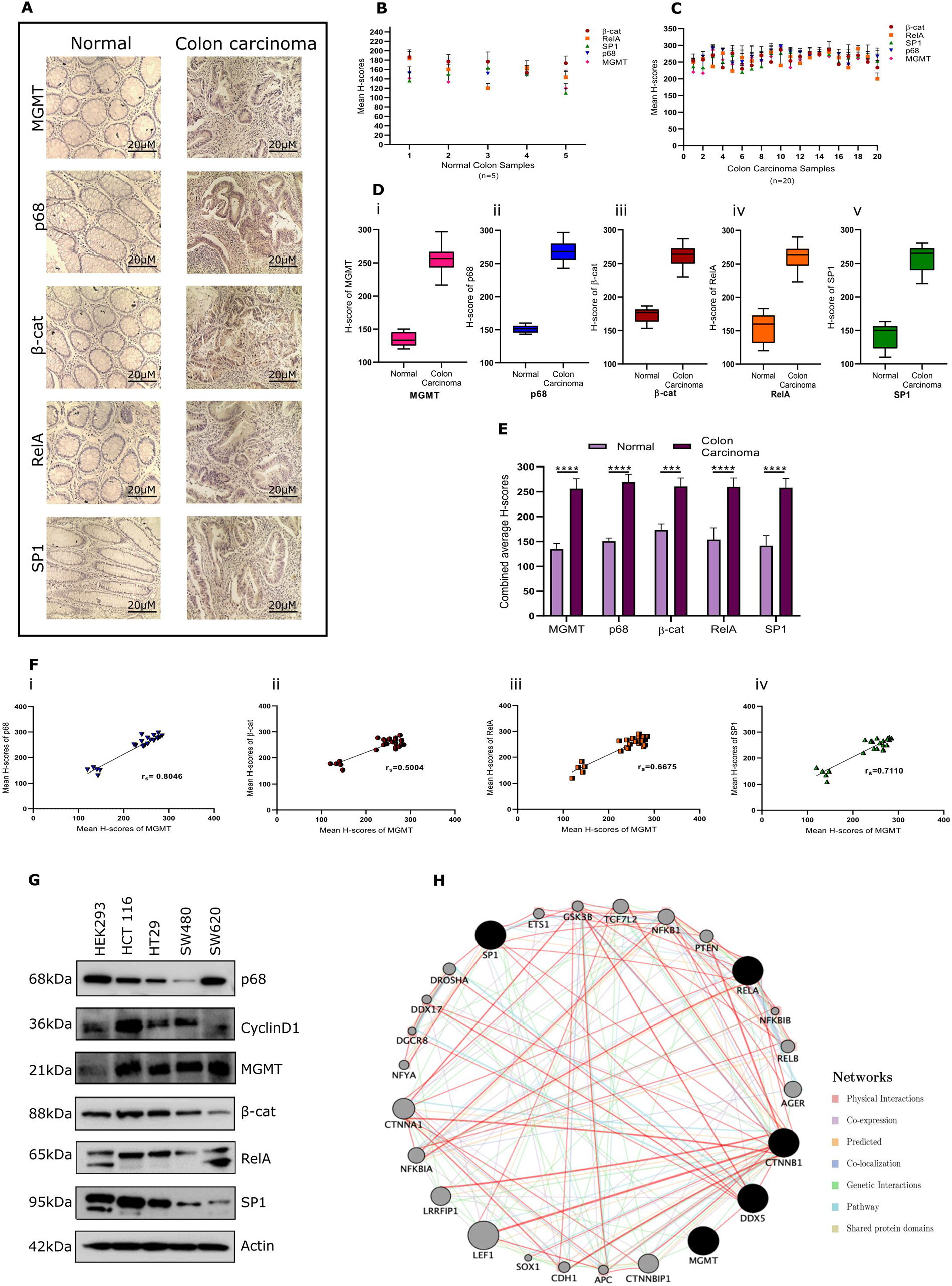
Concomitant gain of MGMT and p68 along with β-catenin, RelA, SP1 gene expressions as they share a strong positive correlations in human colon carcer. **(A)** Representative IHC images of the candidate proteins in human colon carcinoma and adjacent normal colon tissue samples. All images are taken at ×20 magnification. Scatter-plot representation of the mean H-scores of MGMT, p68, β-catenin, RelA and SP1 in **(B)** adjacent normal colon (n = 05) and **(C)** colon carcinoma (n = 20) **(D)** Box-plots depicting distribution of H-scores of MGMT, p68, β-catenin, RelA and SP1 in normal colon and colon carcinoma samples as obtained through calculations from M–W U-test. **(E)** Graphical representation of the combined average H-scores of MGMT, p68, β-catenin, RelA and SP1. Error bars represent the mean (+) s.d.; P < 0.0001 is represented as ****; calculated using Student’s t-test **(F)** Depiction of correlation coefficient (rs) between mean H-scores of (i) MGMT and p68, (ii) MGMT and β-catenin, (iii) MGMT and RelA and (iv) MGMT and SP1 as estimated from IHC images of both normal colon and colon carcinoma tissues combined. **(G)** Immunoblot shows differential expression of p68 and MGMT in multiple colon cancer cell lines. Three out of four colon cancer cell lines as depicted in the subsequent figures were used in this study. **(H)** A GeneMANIA interaction network of DDX5, β-catenin, RelA, SP1 and MGMT was visualized with Cytoscape showing interaction strength by edge thickness, interaction type by color, different edges between nodes and protein score based on node size.

Next, overall expression profiles of MGMT and DDX5 in various cancers were generated using UALCAN by analysis of RNA expression data in 24 different cancer types available in The Cancer Genome Atlas (TCGA) database.^20^ The differential gene expression profiles of MGMT and DDX5 represented their upregulation in most cancer types as compared to their respective normal samples [Supplementary figures S2A & B]. From these cancer types, Colorectal Adenocarcinoma (COAD) was chosen for further analysis. Therefore, we assessed stage specific variations of MGMT and DDX5 expressions in Colorectal Adenocarcinoma (COAD) represented as box plots [Supplementary figures S2C & D]. An increase in MGMT and p68 expression levels was observed with increasing stages of colorectal carcinoma progression vs. normal samples.

### p68 regulates the expression and promoter activity of MGMT gene

To investigate and evaluate the possible mechanism of MGMT gene expression in colon cancer, we initially checked the endogenous expression of MGMT in HCT 116 cells by immuno-cytochemical analysis. We observed p68 is diffusely expressed all over the nucleus and cytoplasm, while interestingly, upon p68 overexpression, it significantly accumulates in numerous nuclear foci visible through GFP-p68 expression whereas upon p68 depletion a decline in overall p68 level was observed. Simultaneously, we observed a visible increase in MGMT expression (RFP-tagged) specifically in those cells that show an enhanced expression of p68 whereas p68 depletion corroborates with decline in MGMT expression (Fig. 2A (i & ii)).

**Fig. 2.**
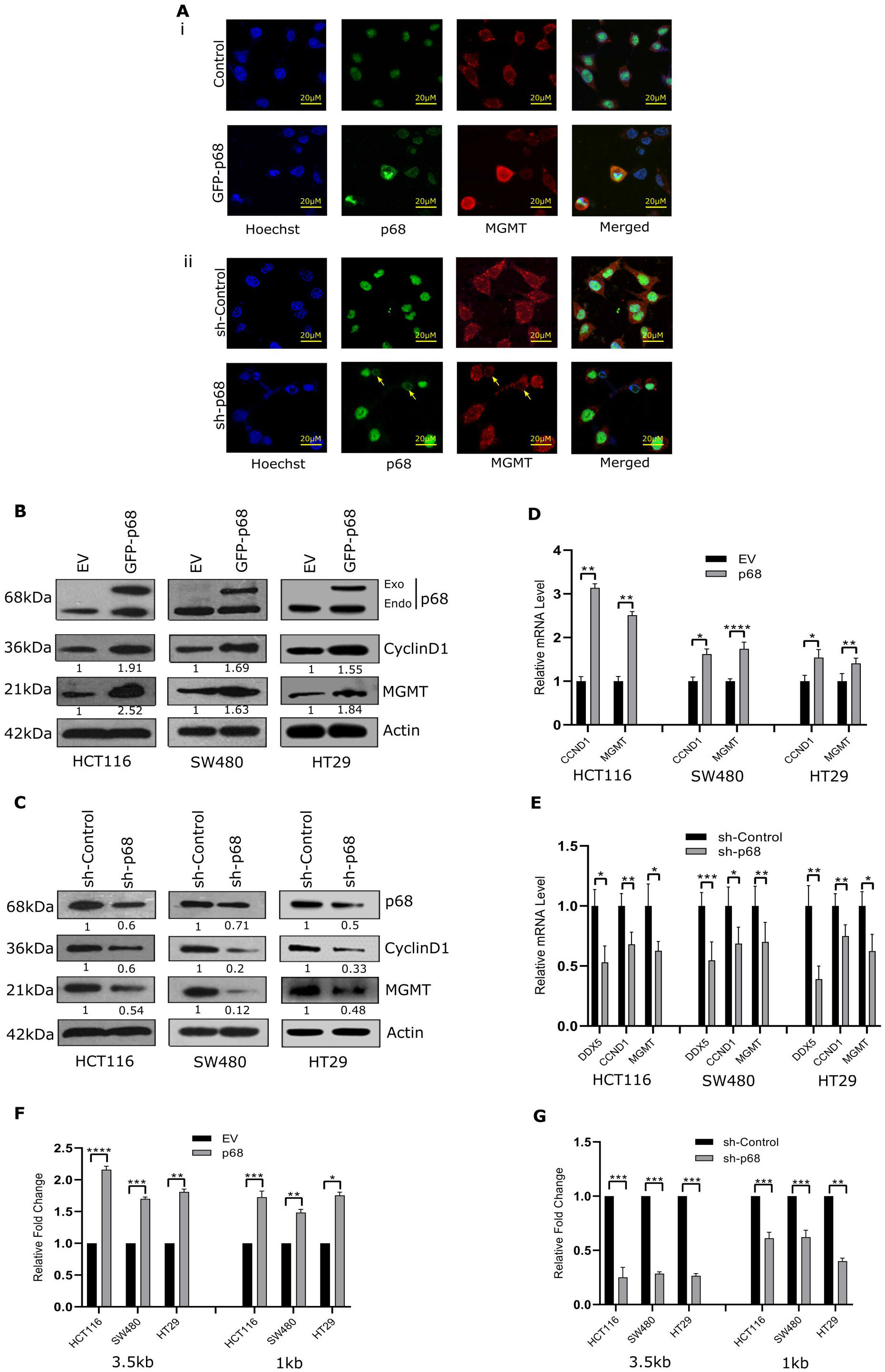
P68 positively regulates MGMT expression and promoter activity in colon cancer cell lines. **(A)** Expression of p68 and MGMT in HCT 116 cells transfected with (i) GFP-p68 or empty vector for 48 hr and (ii) either control shRNA or shRNA against p68, then detected by immuno-cytochemistry. All images were taken at ×20 maginification. Scale bars, 20 μm. Immunoblot analysis of cells transfected with **(B)** either GFP-p68 or the empty vector and harvested at a total of 48 hr post transfection and **(C)** either control shRNA or shRNA against p68 and harvested 48 hr post transfection. Relative mRNA expression (w.r.t 18s rRNA) of MGMT assessed in HCT 116, SW 480 and HT29 cells **(D)** qRT-PCR analysis of cells transfected with GFP-p68 and **(E)** qRT-PCR analysis of cells transfected with either control shRNA or shRNA against p68. Luciferase activity of MGMT promoters (3.5kb and 1kb variants) in colon cancer cell lines co-transfected with **(F)** pIRES-p68, pGL2-MGMT-prom, and pRL-TK (Renilla luciferase construct) for 48 hr. The figure represents relative fold change in luciferase readings, normalized against Renilla reporter activity, measured at 48 hr post transfection and **(G)** either control shRNA or shRNA against p68, pGL2-MGMT-prom and pRL-TK, measured 48 hr after transfection. All experiments were performed in triplicates, n =3. Error bars represent the mean (±) S.D. of independent two-tailed Student’s t-tests, where P<0.0001 is represented as **** for highly significant.

Next, we overexpressed p68 in multiple colon cancer cell lines and the immunoblotting results indicated an enhanced expression of MGMT in all the cases. CyclinD1 is a reported target gene of p68 action^21^ and Actin serves as loading control (Fig. 2B). To study whether the converse holds true, we depleted endogenous p68 in these cell lines by using shRNA against p68 and observed a sharp decline in MGMT expression along with that of Cyclin D1 (Fig. 2C). Further to study whether the effect of p68 on MGMT gene expression is due to its transcriptional regulation, we quantified its mRNA expression and found increase in MGMT mRNA expression upon overexpressing p68 (Fig. 2D), whereas p68 depletion led to a significant reduction in MGMT mRNA level (Fig. 2E). Altogether, we observed that cellular p68 level correlates with MGMT expression in human colon cancer cell lines and p68 positively regulates its expression *in vitro* as a transcriptional co-activator. This led us to explore whether p68 controls MGMT promoter activity. We went on to work with two MGMT promoters of different lengths of 3.5 kb and ∼1kb (precisely 954bp upstream) in HCT116, SW480, HT29 cell lines to better understand the involvement of p68. We observed that p68 overexpression led to an increase in MGMT luciferase activity (Fig. 2F). The reverse was also true when endogenous p68 depleted with shRNA against it that led to a decline in MGMT luciferase activity (Fig. 2G). Moreover, no distinct difference was observed between the two promoter variants. Taken together, these results imply that p68 regulates MGMT promoter activity and its expression in multiple colon cancer cell lines.

### MGMT expression and promoter activity are mediated through β–catenin, RelA and SP1

DDX5 (p68) binds to its target promoters *via* interaction with transcription factors of multiple signalling pathways. Since p68 cannot interact directly with the MGMT promoter region, we investigated the possible involvement of known transcription factors *viz*., β-catenin, RelA and SP1 in colon cancer cell lines. Upon individual overexpression of GFP-tagged β-catenin, RelA and SP1 individually in HCT 116 cells followed by ICC (immuno-cytochemical) staining, we observed elevated expression of MGMT (RFP-tagged) specifically in those cells that show enhanced expression of β-catenin, RelA and SP1 (Fig. 3A (i), B (i) & C (i)). Next, we were intrigued to study the effect on MGMT expression by depletion of endogenous levels of β-catenin, RelA and SP1 individually by their respective inhibitors, and observed overall decline in their expression levels along with a marked decrease in MGMT expression (Fig. 3A(ii), B(ii) & C(ii)). Here, FH535, Bay11 and Mithramycin A are the antagonistic inhibitors of β-catenin, RelA and SP1 respectively and were used at a concentration of 15μM, 10μM and 24nM correspondingly for 24 hr. We then immediately decided to study MGMT promoter activity, hence overexpressed as well as downregulated β-catenin, RelA and SP1 individually in HCT 116 and SW480 colon cancer cell lines. The effect of individual overexpression of each of the three transcription factors corroborated to an increase in MGMT promoter activity (Fig. 3D (i, ii & iii)) and the effect of individual depletions of each of the three transcription factors using their respective shRNAs corroborated to decrease in MGMT promoter activity (Fig. 3E (i, ii & iii)). Hence, β-catenin, RelA and SP1 pathways might be responsible for triggering and positively regulating MGMT gene expression. The luciferase activity remained unchanged by their empty vector control, as was expected.

**Fig. 3.**
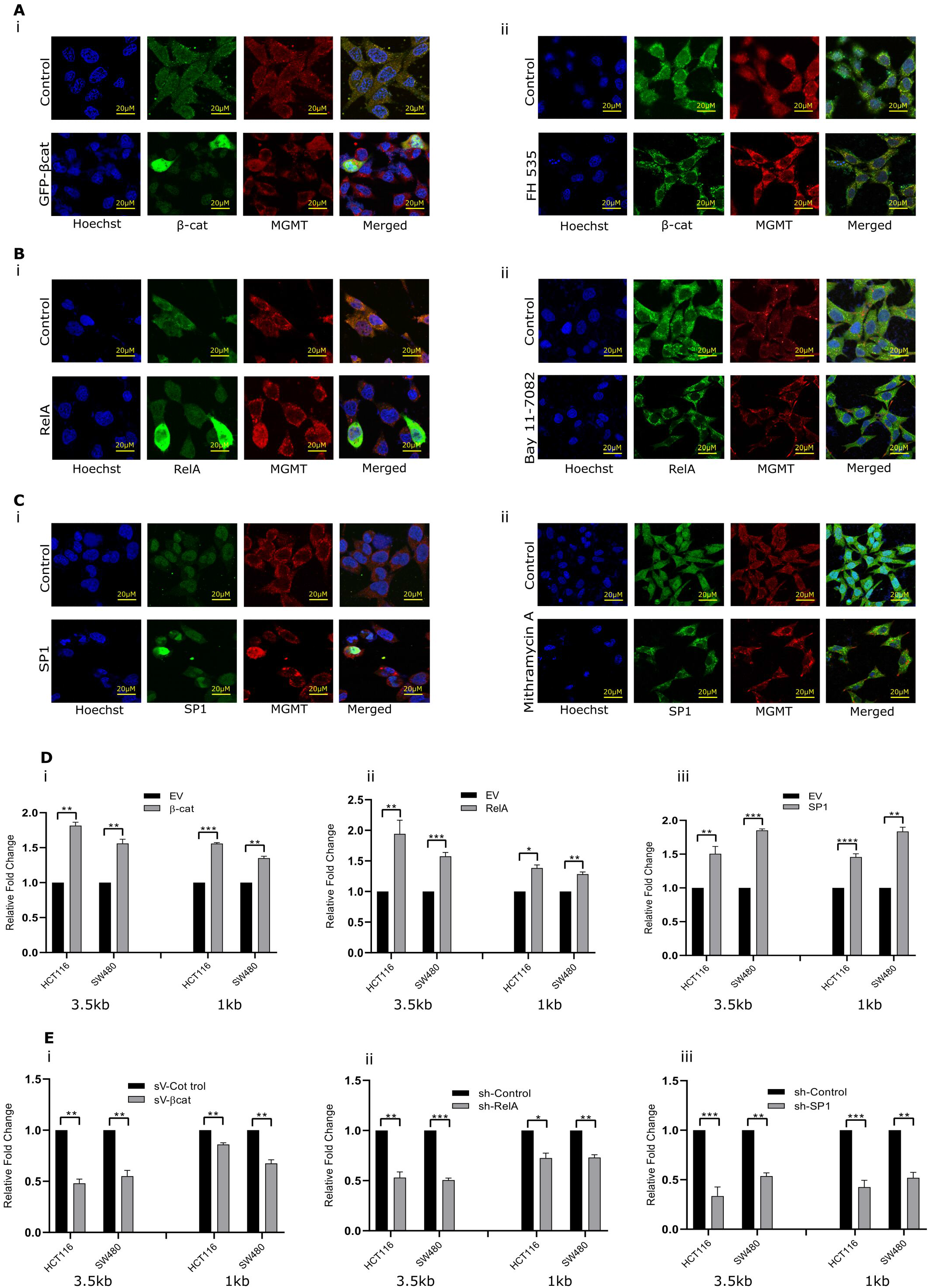
MGMT expression and promoter activity is mediated by β-catenin, RelA and SP1. Expression of MGMT (RFP-tagged secondary antibody was used) with respect to β-catenin, RelA and SP1 in HCT 116 cells transfected with either empty vector or **(A** (i)) GFP-β-catenin vector, **(B** (i)) pCMV4-RelA (GFP-tagged secondary antibody was used) and **(C** (i)) pcDNA3.1-SP1 (GFP-tagged secondary antibody was used) for 48 hr and then detected by immuno-cytochemistry. Expression of MGMT (RFP-tagged secondary antibody was used) with respect to β-catenin, RelA and SP1 in HCT 116 cells treated with either DMSO or **(A** (ii)) FH535 (15μM), **(B** (ii)) Bay11 (10μM) and **(C** (ii)) Mithramycin A (24nM) (GFP-tagged secondary antibody was used for each of them) for 24 hr and then detected by immuno-cytochemistry. All images were taken at ×20 maginification. Scale bars, 20 μm. Luciferase activity measured in cells co-transfected with **(D)** either control vector or (i) β-catenin, (ii) RelA and (iii) SP1 individually and **(E)** either control shRNA or shRNA against (i) β-catenin, (ii) RelA and (iii) SP1 individually, pGL2-MGMT-prom (3.5kb and 1kb variants) and Renilla luciferase construct pRL-TK (50ng) for 48 hr. The figures represent relative fold change in luciferase readings, normalized against Renilla reporter activity. All experiments were performed in triplicates, n =3. Error bars represent the mean (±) S.D. of independent two-tailed Student’s t-tests, where P<0.0001 is represented as **** for highly significant.

### β–catenin, RelA and SP1 positively regulate MGMT mRNA and protein levels

To examine *in vitro* correlations of β-catenin, RelA and SP1 with MGMT, we individually overexpressed them in HCT 116, SW480 and HT29 cell lines and found enhanced expression of MGMT in all the cases (Fig. 4A, 4E & 4I). Whereas, upon individual downregulation of β-catenin, RelA and SP1 in these cell lines by using their respective shRNAs against them, we observed decreased MGMT expression. Here, we kept Cyclin D1 and XIAP as positive controls (Fig. 4B, 4F & 4J). CyclinD1 is a reported target gene of β-catenin^22^ whereas XIAP has been reported as target gene of both RelA and SP1^23, 24^ and Actin serves as loading control. Then, to study whether the effect of any of these transcription factors *viz.,* β-catenin, RelA and SP1 on MGMT gene expression is due to its transcriptional regulation, we quantified their mRNA expressions and found increased expression of MGMT mRNA upon their individual overexpression (Fig. 4C, 4G & 4K). We also observed that individual depletion of β-catenin, RelA and SP1 by their respective shRNAs led to significant decrease in MGMT mRNA levels (Fig. 4D, 4H & 4L). Taken together, these results imply that β-catenin, RelA and SP1 positively regulate both protein and mRNA expressions of MGMT in multiple colon cancer cell lines. The above findings were further verified in HT29 cell line and represented in Supplementary figure S3.

**Fig. 4.**
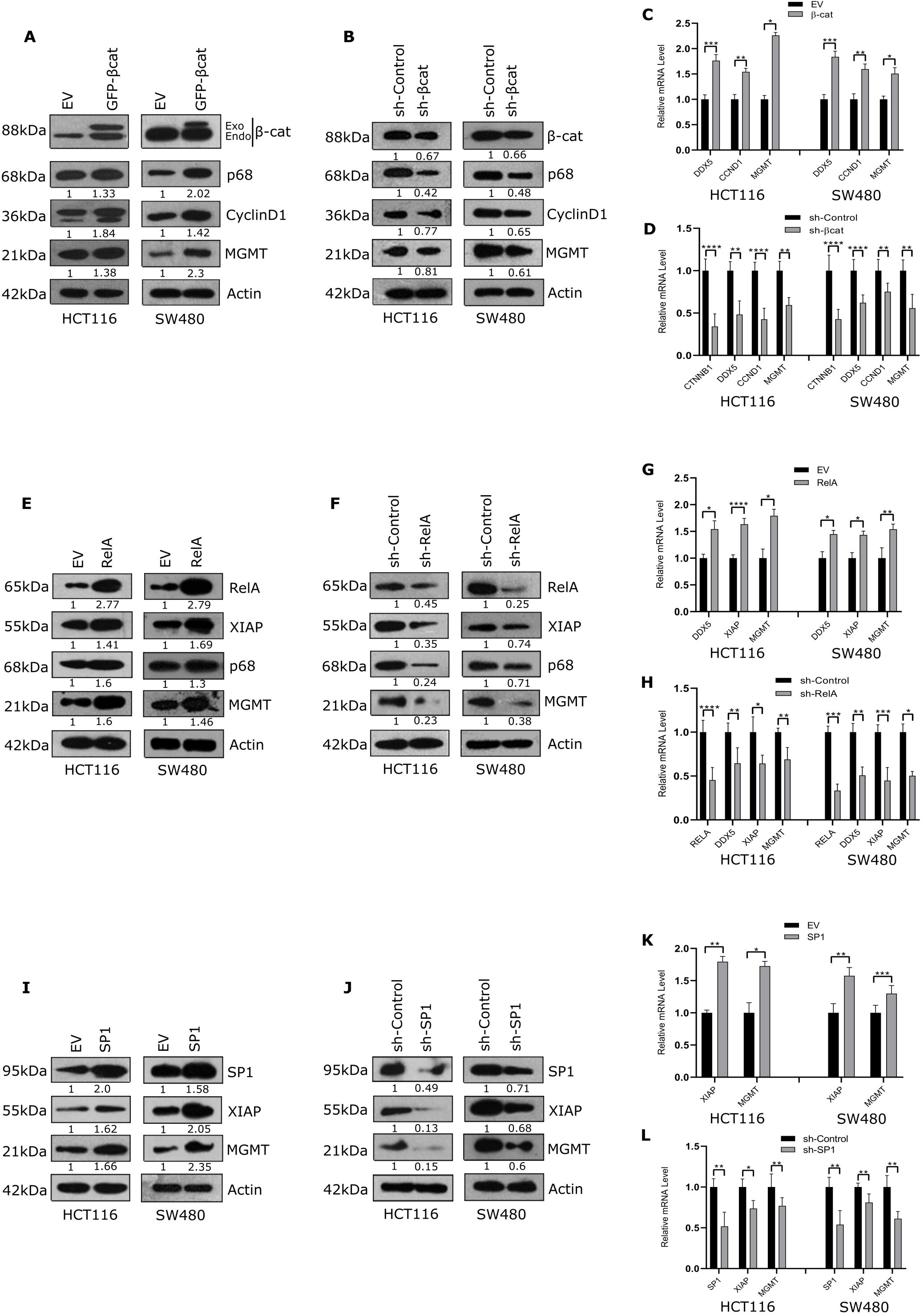
β-catenin, RelA and SP1 positively regulate MGMT mRNA and protein levels. Immunoblot analysis of colon cancer cells transfected with either **(A)** GFP-β-catenin or the empty vector, **(E)** pCMV4-RelA or the empty vector and **(I)** pcDNA3.1-SP1 or the empty vector were harvested at 48 hr post transfection. Immunoblot analysis of cells transfected with either control shRNA or shRNA against **(B)** β-catenin **(F)** RelA and **(J)** SP1 were harvested 48 hr post transfection. Relative MGMT mRNA expression (w.r.t 18s rRNA) was assessed in HCT 116 and SW480 cell lines. qRT-PCR analysis of cells transfected with **(C)** GFP-β-catenin, **(G)** pCMV4-RelA and **(K)** pcDNA3.1-SP1. qRT-PCR analysis of cells transfected with either control shRNA or shRNA against **(D)** β-catenin, **(H)** RelA and **(L)** SP1 and harvested 48 hr post transfection. Error bars represent the mean (±) S.D. of independent two-tailed Student’s t-tests, where P<0.0001 is represented as **** for highly significant.

### P68 colocalizes and interacts with SP1

Overexpression of p68 in multiple cancers including colon cancer has been substantiated by several studies. The role of p68 as a co-activator of transcription factors β-catenin and NF-κB (p50-p65) has been previously explored by us and others.^16, 21^ We were interested to ascertain these interactions in colon cancer cell line, hence, a co-localization experiment was conducted in HCT 116 cells to explore the possible interaction of p68 with β-catenin and RelA independently, and a strong colocalization of β-catenin and p68 as well as RelA and p68 was detected by ICC (Fig. 5A). In addition to involvement of β-catenin and RelA, in regulating MGMT gene expression, we were intrigued to explore the colocalization and physical interaction between p68 and SP1. Hence, we conducted a co-localization study in HCT116 and HEK293 through immunofluorescence analysis. As expected, p68 and SP1 significantly colocalized in both the cell lines (Fig. 5B). Thus, these colocalization data indicated their possible interaction across normal and cancerous scenarios

**Fig. 5.**
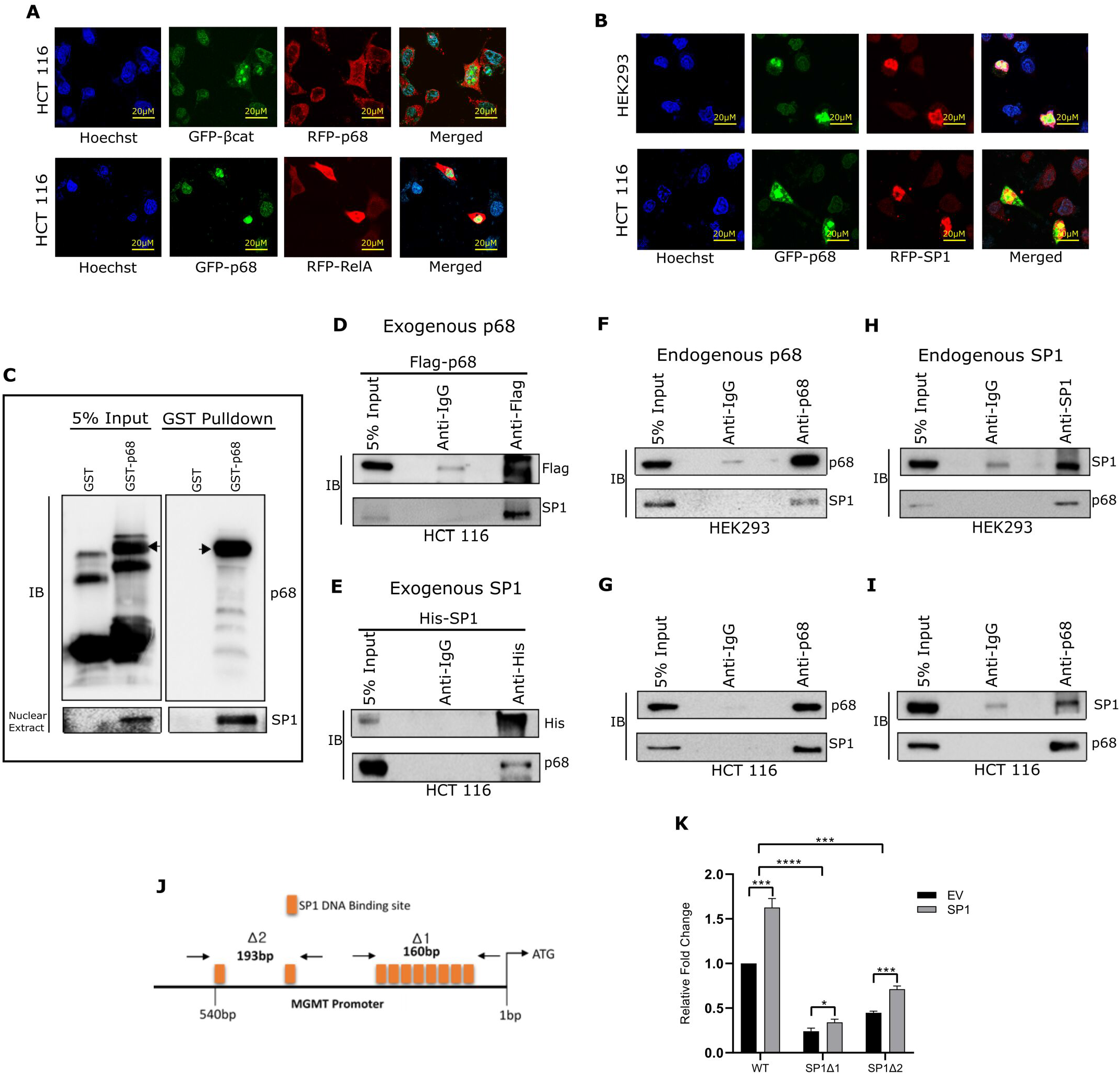
p68 co-localizes and physically interacts with SP1. Expression of p68 and **(A)** β-catenin or RelA in HCT 116 cells respectively **(B)** SP1 in HCT 116 and HEK293 cells as detected by immuno-cytochemistry. All images were taken at ×20 magnification. Scale bars, 20μm. All experiments were performed, n=3. GFP-β-catenin and Flag-p68 were co-transfected where RFP-tagged secondary antibody was used for p68 detection. Similarly, GFP-p68 was co-transfected with pcMV4-RelA or pcDNA3.1-SP1 individually and were visualized by RFP-tagged secondary antibody for RelA and SP1. (**C)** Purified GST-tagged p68 or GST were incubated with whole cell extracts of HEK293 cells grown in 100 mm cell culture dishes and transfected with 8 μg of pcDNA3.1-SP1. This was followed by pull down with GST beads and analysis of the indicated proteins by immunoblotting. 5% of the whole cell extract and purified proteins were used as input; specific bands are indicated by arrows. HCT 116 cells were grown in 100 mm cell culture dishes and were transfected with 4 μg of pIRES-p68 or 4 μg of pcDNA3.1-SP1. 48 hr post-transfection, whole cell extracts were prepared followed by IP with **(D)** anti-Flag and **(E)** anti-His antibodies and analysis of the indicated proteins by immunoblotting with the protein tag specific antibody (Flag for p68 and His for SP1). IP with normal rabbit immunoglobulin G (IgG) served as negative control. Whole cell extracts were prepared from HEK293 and HCT 116 cells, respectively, followed by IP with **(F & G)** anti-p68 or **(H & I)** anti-SP1 and analysis of the indicated proteins by immunoblotting. IP with normal rabbit IgG served as negative control. In all, 5% of the whole cell extract was used as input. **(J)** Schematic representation of wild type MGMT promoter (1 kb variant) mentioning all its SP1 binding domains and of the mutant MGMT promoter constructs pGL2-MGMT-prom (Δ1 40-200, Δ2 347-540), with its SP1 binding domains deleted. **(K)** Luciferase activity of HCT 116 cells transfected with either WT pGL2-MGMT-prom (1kb variant) or its mutant variants (Δ140-200 or Δ2 347-540) in SP1 overexpression system along with pRL-TK, measured 48 hr after transfection.

Next, we investigated physical interaction between p68 and SP1. First, glutathione S-transferase (GST) pull-down assays were performed with purified GST or GST-p68 and whole cell lysates of HEK293 cells overexpressing SP1. GST-p68 interacted with SP1 thereby suggesting physical interaction between them (Fig. 5C). Additionally, immunoprecipitation (IP) assay using whole cell lysates of HEK293 cells showed that exogenously expressed p68 distinctly co-precipitated with exogenously expressed SP1 and vice versa (Supplementary figure S4). Furthermore, IP assay using whole cell lysates of HCT 116 cells showed endogenous SP1 interacts with exogenously expressed p68 and vice versa (Fig. 5D & 5E). Moreover, IP assays were conducted using whole cell lysates of both HCT116 and HEK293 cells where endogenous SP1 was readily found in the immuno-complex of endogenous p68 and vice versa (Fig. 5F, 5G, 5H & 5I). Thus, all these pulldown studies and immunoprecipitation results strongly suggest physical association and interaction of p68 and SP1 in both normal and cancer cells.

Further, we analysed the promoter regions of the human MGMT gene as retrieved from both Alggen PROMO database and Eukaryotic Promoter database. From MGMT promoter study, we found 10 very closely spaced SP1 binding sites within 550 bps upstream of +1 site. To better understand the involvement of SP1 sites on MGMT promoter activity we generated two SP1 mutant promoter constructs [Δ1 40-200 (includes 8 sites), Δ2 347-540 (includes 2sites)] of the MGMT 1kb promoter and introduced them individually or WT-MGMT promoter in HCT 116 cells to study MGMT promoter activity (Fig. 5J). As expected, both the mutants were less active than the WT and the Δ1 mutant showed lesser MGMT promoter activity than the Δ2 mutant, indicating that all SP1 sites are important for MGMT gene expression (Fig. 5K).

### P68 acts as a master regulator of MGMT gene expression mediated through β-catenin, RelA and SP1

MGMT promoter contains seven TCF-4/LEF (for β-catenin), seven RelA and ten SP1 binding sites in a stretch of -3500 to +24 bp. Hence, we delved deeper into the study of promoter occupancy by chromatin immunoprecipitation (ChIP) assay. We designed ChIP specific primer sets for each of β-catenin, RelA and SP1 based on their binding sites as depicted by schematic diagrams Fig. 6A (i, ii & iii).

**Fig. 6.**
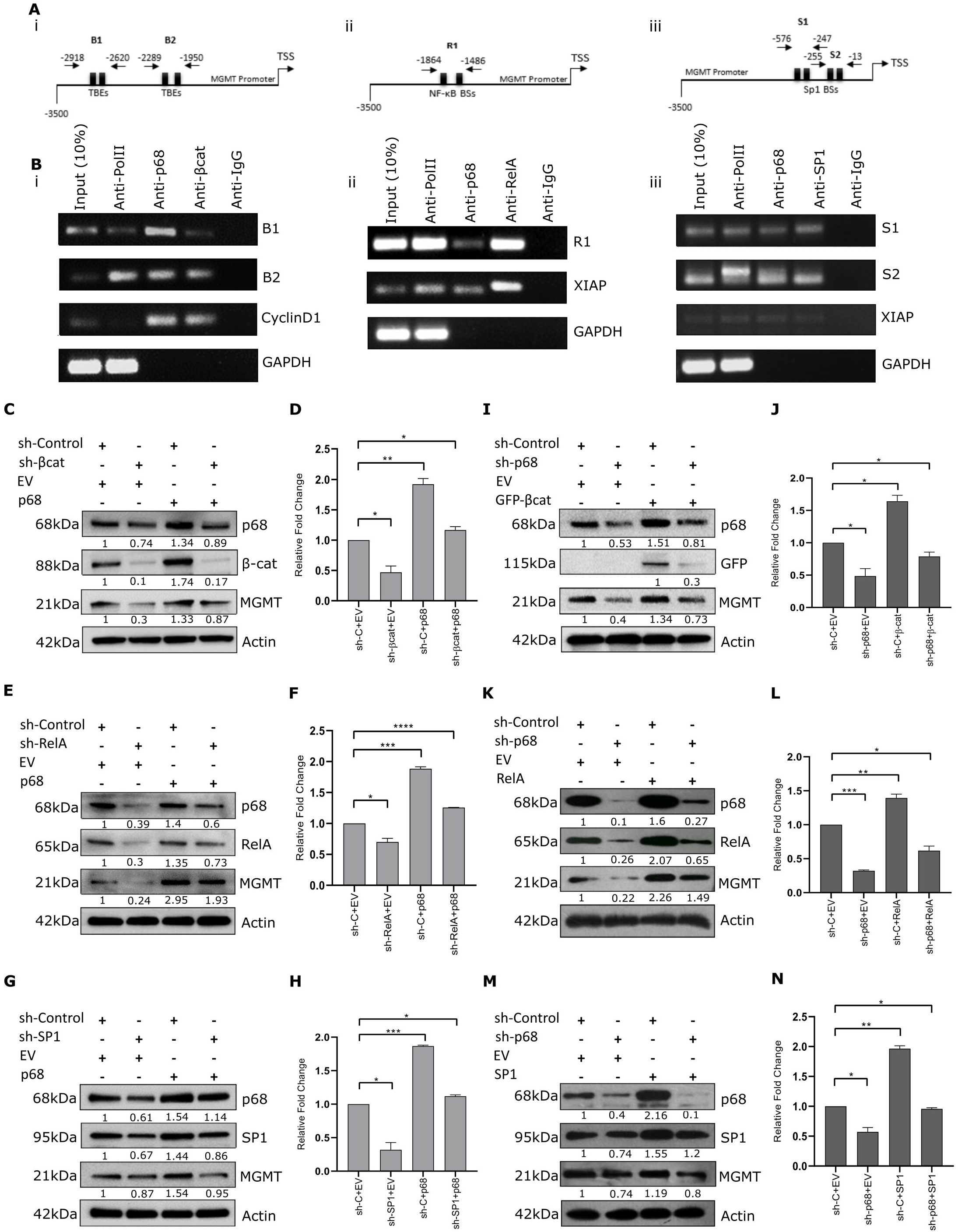
p68 associates with the MGMT promoter and acts as a master regulator of β-catenin, RelA and SP1 mediated MGMT gene expression. **(A)** Schematic representation of human (-3500 to TSS) MGMT promoter with ChIP primers designed against (i) TCF-4/β-catenin binding sites (ii) NF-κB/RelA binding sites (iii)SP1 binding sites. **(B)** Chromatin immunoprecipitation (ChIP) assay performed in human HCT 116 cell line targeting endogenous MGMT promoter and respective positive controls using (i) p68 and β-catenin antibodies, (ii) p68 and Rela antibodies and (iii) p68 and SP1 antibodies. IgG antibody was used as negative control in all cases. HCT 116 cells were transfected with shRNA targeting **(C)** β-catenin, **(E)** RelA and **(G)** SP1 or control shRNA independently for 24 hr, with subsequent overexpression of p68 for another 24 hr and expression of respective protiens were analysed by IB after 48 hr of total transfection. Luciferase activity of MGMT promoter co-transfected with control shRNA or shRNA against **(D)** β-catenin, **(F)** RelA or **(H)** SP1 individually in separate sets, pGL2-MGMT-prom and pRL-TK and then 24 hr later transfected with pIRES-p68 individually and measured 48 hr after transfection. HCT 116 cells were transfected with small shRNA targeting p68 or control shRNA. After 24 hr, cells were transfected with **(I)** GFP-β-catenin and kept for another 24 hr. Whole cell lysates (WCLs) were prepared and proteins were analysed by immunoblotting (IB) as shown in the figure. P68 was knocked down followed by overexpression of **(K)** RelA or **(M)** SP1 by ectopic expression of both of them respectively in separate sets in HCT 116 cells. WCLs of 48 hr post-transfected cells were analysed for MGMT expression by IB. Luciferase activity of MGMT promoter co-transfected with control shRNA or shRNA against p68, pGL2-MGMT-prom and pRL-TK and then 24 hr later transfected with **(J)** β-catenin, **(L)** RelA or **(N)** SP1 individually in separate sets and measured 48 hr after transfection. Error bars represent the mean (±) S.D. of independent two-tailed Student’s t-tests, where P<0.0001 is represented as **** for highly significant.

The ChIP assays on endogenous MGMT promoter were performed with cross-linked chromatin fragments prepared from HCT 116 cells using respective antibodies as depicted in the figure to elucidate the involvement of: (i) p68 and β-catenin, (ii) p68 and Rela, and (iii) p68 and SP1 on MGMT promoter (Fig. 6B (i, ii & iii)). RNA polymerase II and anti-IgG antibodies were chosen as positive and negative controls, respectively. We observed that β-catenin occupies the MGMT promoter as well as the Cyclin D1 promoter (positive control). P68 specific enrichment of β-catenin sites was found on MGMT promoter (Fig. 6B (i)). Similarly, we found enrichment of RelA and SP1 on MGMT promoter as well as XIAP promoter (positive control). P68 showed enrichment at both RelA and SP1 sites on MGMT promoter as well as of XIAP promoter (indicating potential p68 involvement in XIAP regulation) (Fig. 6B (ii & iii)). The ChIP data using chromatin of HCT 116 cells demonstrated that p68 binds to MGMT promoter through β-catenin, RelA and SP1 binding sites.

Furthermore, to decipher the significance of p68 in the β-catenin, RelA and SP1-mediated MGMT gene expression, we individually knocked down β-catenin, RelA and SP1 using their corresponding shRNA in HCT 116 cells for 24 hr, followed by transiently overexpressing p68 for another 24 hr. We observed increased expression of MGMT by p68 without individual additional signalling by β-catenin, RelA or SP1 expression (Fig. 6C, 6E & 6G). P68 could itself rescue MGMT expression even in the knocked down condition of β-catenin, RelA or SP1. Surprisingly, here we also observed that in SP1 knockdown condition, the ability of p68 to rescue MGMT expression is comparatively low as compared to the knockdown condition of β-catenin or RelA. These results clearly indicate that SP1 is a strong regulator of MGMT gene expression further enhanced by p68. To further establish the involvement of p68 in MGMT expression through transcriptional activation of its promoter, we tried to rescue the decreased MGMT promoter activity by p68 overexpression in the individual knockdown condition of β-catenin, RelA or SP1. We chose the 3.5 kb variant of MGMT promoter because it has multiple β-catenin, RelA and SP1 binding sites, thus would help us to better understand the involvement of p68 on MGMT gene regulation. Luciferase assays with cells 48 hr post-transfection indicated an increase in MGMT promoter activity by p68 as compared to the empty vector controls (Fig. 6D, 6F & 6H).

Next, we also decided to analyse MGMT protein expression under p68 downregulated condition followed by upregulation of β-catenin, RelA or SP1 individually. This regulation was confirmed by knocking down p68 in HCT 116 cells for 24hr followed by β-catenin, RelA or SP1 overexpression individually for the next 24 hr. We observed that p68 downregulation has a severe effect on the resultant β-catenin, RelA or SP1-dependent expression of MGMT (Fig. 6I, 6K & 6M). We also explored the effect of individual upregulation of β-catenin, RelA and SP1 pathways under p68 knocked-down condition on MGMT promoter activity. MGMT promoter showed a decline in its activity when p68 was downregulated as compared to the empty vector control (Fig. 6J, 6L & 6N). These results, in turn, establishes the fact that p68 truly controls MGMT promoter activity as a strong co-activator of β-catenin, RelA and SP1-mediated signalling, thus acting as a master regulator.

The importance of MGMT in conferring chemoresistance to DNA-alkylating drugs, the most prominent being TMZ, in patients across cancers is well established. Thus, we were intrigued to further explore the involvement of p68 as a master regulator of MGMT in manipulating TMZ based response in human colon cancer cell lines.

### p68–‘β-catenin/RelA/SP1’–MGMT circuits are responsible for TMZ chemoresistance

DDX5 (p68) is an established oncogenic transcriptional coactivator playing its role in multiple cancers. To see the involvement of p68–‘β-catenin/RelA/SP1’–MGMT networks on TMZ-induced chemoresistance, we carefully monitored the effect of p68 on TMZ treated cells. For this we first performed Clonogenic assay in p68 overexpressed and knocked down HCT 116 cells. We found an increase in the number and size of the colonies and vice-versa in p68 overexpressed and knocked down conditions, respectively, and represented as bar graphs (Fig. 7A (i & ii)). Further, wound healing assay was performed after p68 overexpression, which showed an increased healing in the wound region after 24 hr of the treatment thus highlighting that p68 upregulation aids in cellular proliferation (Fig. 7B (i)). A considerable decrease in the wound healing was observed in p68 knockdown condition as compared to its control and has been demonstrated in the bar graphs (Fig. 7B (ii)). These results ascertain the ploriferative potential of p68.

**Fig. 7.**
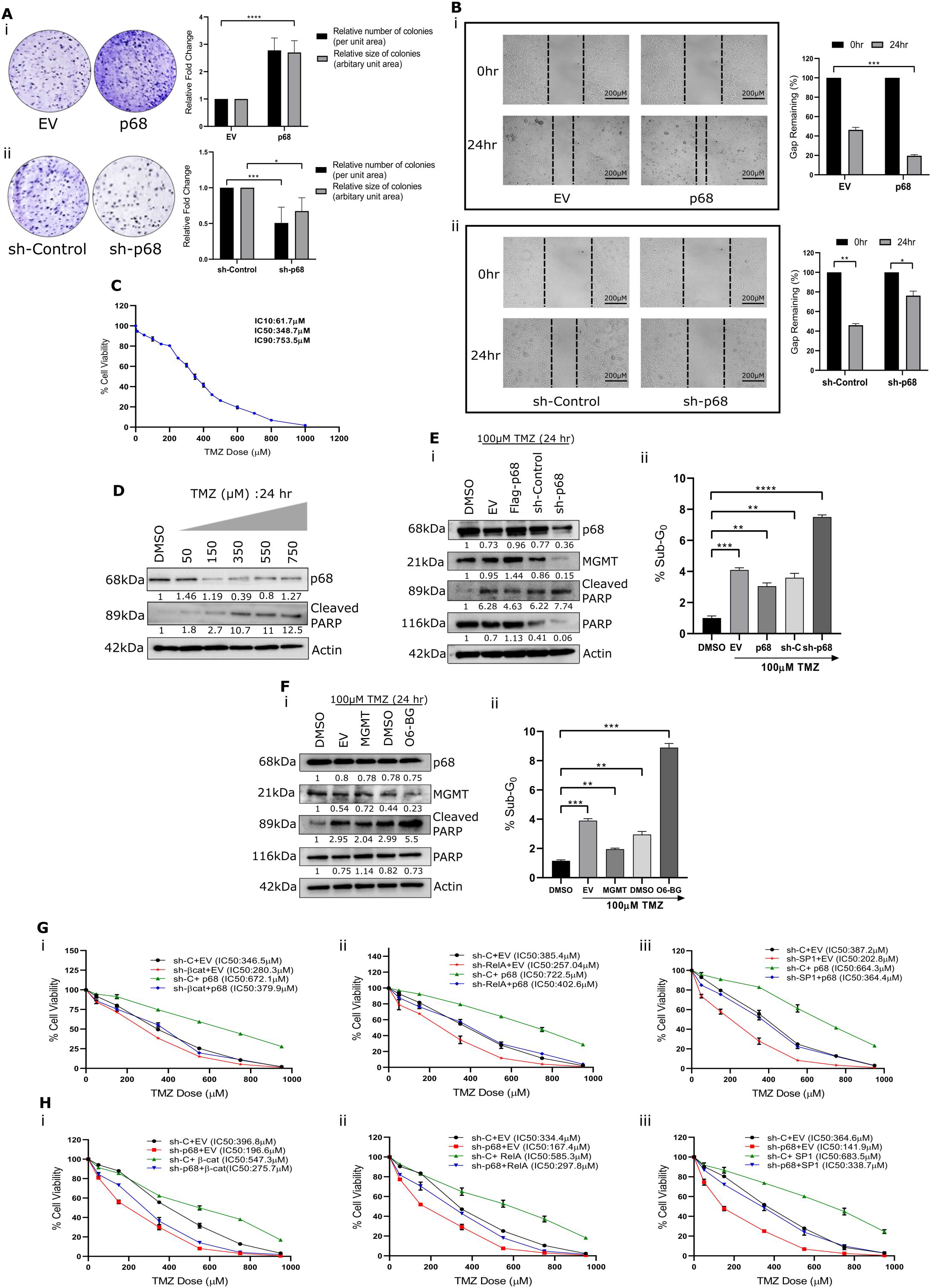
P68 actively regulates MGMT mediated TMZ response in HCT 116 cells. **(A)** Cell proliferation assay measured in HCT 116 cells transfected with (i) either GFP-p68 or empty vector for p68 overexpression and (ii) either shRNA against p68 or control shRNA for p68 knock down and kept for 15 days and as observed for colony formation capacity (n=3). The size and number of colonies are represented in the bar diagram. **(B)** Scratch assay performed on HCT 116 cells subjected to (i) either GFP-p68 or empty vector and (ii) either shRNA against p68 or control shRNA for a period of 24 hr. Scale bar of 200 μm applies to all the images. Percentage of the gap remaining measured is represented as bar diagram. **(C)** IC10 ∼62μM, IC50 ∼349μM and IC90 ∼754μM doses of TMZ in HCT116 cells were determined by MTT Assay for 24 hr. **(D)** Expression pattern of p68 and Cleaved PARP was studied by IB under various doses of TMZ (50μM-750μM) for 24 hr. Actin was used as loading control and DMSO was used as experimental control. **(E)** HCT 116 cells transfected with either pIRES-p68/control empty vector for p68 overexpression or with shRNA against p68/control shRNA for p68 knockdown and then 24 hr later treated with 100μM of TMZ for next 24 hr were then (i) analysed by immune-blotting for desired proteins and (ii) processed for cell cycle analysis by flow cytometry. Graph was plotted depicting the SubG0 percentage (n=3). **(F)** HCT 116 cells transfected with pSV2-MGMT/control empty vector (PGZ) for MGMT overexpression or treatment with 2.5μM O6-BG/DMSO control for MGMT inhibition and then after 24 hr of transfection/treatment, treated with 100μM TMZ for 24 hr were then (i) analysed by immunoblotting for desired proteins and (ii) processed for cell cycle analysis by flow cytometry. Graph was plotted depicting the Sub-G_0_ percentage (n=3). **(G)** HCT 116 cells were transfected with shRNA targeting (i) β-catenin, (ii) RelA and (iii) SP1 or control shRNA independently for 24 hr, with subsequent overexpression of p68 for another 24 hr. TMZ treatment was done with varying doses (10-1000μM) and MTT assay was conducted after 72 hr of total transfection. **(H)** HCT 116 cells were transfected with small shRNA targeting p68 or control shRNA. After 24 hr, cells were transfected with (i) GFP-β-catenin, (ii) RelA or (iii) SP1 by ectopic expression of all three of them respectively in separate sets in HCT 116 cells for 24 hr. TMZ treatment was done with varying doses (10-1000μM) and MTT assay was conducted after 72 hr of total transfection. Error bars represent the mean (±) S.D. of independent two-tailed Student’s t-tests, where P<0.0001 is represented as **** for highly significant.

Further, IC50 of TMZ was determined by conducting MTT assay on 50-60% confluent HCT 116 cells upon treatment with varying doses of TMZ ranging from 10μM to 1200μM for 24 hr, where DMSO was kept as control. The range was selected after multiple trial and error assays and previous reports. It was found that IC50 of TMZ is ∼349μM, whereas IC10 is ∼62μM and IC90 is ∼754μM (Fig. 7C). Next, the expression of p68 and Cleaved-PARP were studied in HCT 116 cells treated with TMZ (dose ranging from 50-750μM) for 24 hr by immunoblotting (Fig. 7D). We observed a potential correlation between p68 expression attributing to apoptotic hindrance. Additionally, we decided to unravel the intricacy of TMZ-induced apoptosis with the involvement of p68 or MGMT using 100uM TMZ (optimized dose) to observe overall apoptosis (Sub-G_0_ population) by Flow cytometry-based Cell Cycle Analysis and check the status of PARP and Cleaved PARP by immunoblotting to understand the individual impact of p68 or MGMT on TMZ treated colon cancer cells. We observed that either p68 or MGMT overexpression causes a decline in apoptosis whereas knocking down p68 or inhibiting MGMT resulted in increased apoptosis and thus, a better response to TMZ treatment (Fig. 7E & 7F). All raw data of FACS analysis are presented in Supplementary figure S5. All these results further reveal the relationship between p68 and MGMT on TMZ response.

Next, to further explore the possible involvement of p68 in MGMT mediated TMZ chemoresistance *via* β-catenin/RelA/SP1 axes, we performed cell viability assays. We individually knocked down β-catenin/RelA/SP1 using their corresponding shRNA in HCT 116 cells for 24 hr, followed by transiently overexpressing p68 for another 24 hr. P68 could itself significantly increased the IC50 value of TMZ even under individual down regulation of β-catenin/RelA/SP1 resulting in better survival of these cells (Fig. 7G (i, ii & iii)). Surprisingly, here we also observed that in SP1 knockdown condition, the ability of p68 to enhance the % cell viability was comparatively low as compared to β-catenin and RelA mediated signalling. This result indicates that p68-SP1 axis could be a strong regulator of MGMT gene expression. We then decided to analyse the effect of TMZ treatment on % cell viability in the p68 knocked down cells followed by individual upregulation of β-catenin, RelA and SP1. We observed that the resultant knockdown of p68 compromises the β-catenin/RelA/SP1 dependent partial rescue (survival) of TMZ treated cells (Fig. 7H (i, ii & iii)).

## Discussion

Several studies and evidences have elucidated the role of MGMT as a prognostic biomarker for GBM patients and also implicated its similar role in some other cancers. MGMT expression has been co-related to TMZ related chemoresistance in glioma. Under the arena of drug repurposing, TMZ is also being used to treat colon adenocarcinoma and breast cancer. But, here too, chemoresistance remains a challenge. Considering the literature studies, we decided to explore the regulation of MGMT gene in colon cancer. Parallelly, since decades, p68 has been increasingly described as an enzyme closely associated with tumorigenesis. Therefore, we understand the importance of p68 expression in cancer cells, which could further act as a co-activator for multiple transcription factors and regulate a diaspora of cellular events. In this context, the possible role of p68 in regulating MGMT and subsequently, MGMT-related TMZ chemoresistance can be studied in colon cancer.

On pursuing our objective, we observed through relative protein expression patterns that both p68 and MGMT maintain a strong positive correlation in human colon carcinoma and adjacent normal tissues as well as cell lines, with an evidently strong expression in colon carcinoma cells. This correlation hints towards a possible association between these two tumorigenic factors and we explored the signalling cascade that connects them. We observed that p68 enhances MGMT protein and mRNA levels in multiple colon cancer cell lines.

Literature study has revealed that p68 doesn’t possess any DNA binding domain, hence, it is dependent on other transcription factors to regulate its target genes through co-activation. β-catenin, RelA and SP1 are considered as major regulators of MGMT gene, SP1 could be the strongest. Thus, the individual impact of each transcription factor *viz*., β-catenin, RelA and SP1 on MGMT regulation was studied. Our luciferase results complied with our observations, where we found p68-mediated activation of MGMT promoter activity. Two types of MGMT promoters, 3.5kb and 1kb, were chosen to better understand the results. We observed from our experimental data that β-catenin, RelA and SP1 classically activate MGMT transcription by binding through their consensus sites on the MGMT promoter. Since p68 is a co-transcriptional regulator, this further intrigued us to investigate its direct interaction with SP1. We found interaction between p68 and SP1 from IP and pull down assays. Therefore, we proved here a novel interaction between p68 and SP1 (Fig. 5). Thus, SP1 can now also be considered as one of the transcription factors that is co-activated by p68. Additionally, Chromatin immunoprecipitation studies conducted on HCT 116 cell line indicated a strong binding of p68 to β-catenin, RelA and SP1 which bind to their respective sites on MGMT promoter. Through rescue experiments, we observed the importance of p68 in regulating MGMT gene expression. β-catenin, RelA or SP1-mediated signalling cascades optimally function in regulating MGMT in the presence of p68. Furthermore, it was important to understand the possible implications of p68-MGMT signalling network established here.

It was interesting to decipher the involvement of p68 in conferring TMZ-based chemoresistance in colon cancer. Low dose of TMZ, that is, 100μM was chosen to better understand the intricacy in expression patterns. Cell cycle analysis by flow cytometry helped us to understand the intricate dynamics between p68 expression and TMZ-mediated apoptosis. P68 overexpression compromises response to TMZ in HCT 116 cells even within 24 hr, further supporting the involvement p68 in conferring chemoresistance. Additionally, cell viability assays confirmed the role of p68 in manipulating TMZ-induced apoptosis *via* β-catenin/RelA/SP1 axis. Thus in this study, we successfully established p68 as a master regulator of MGMT gene expression and a crucial factor in TMZ chemoresistance mediated through multiple axes like β-catenin, RelA and SP1 (Fig. 8D).

**Fig. 8.**
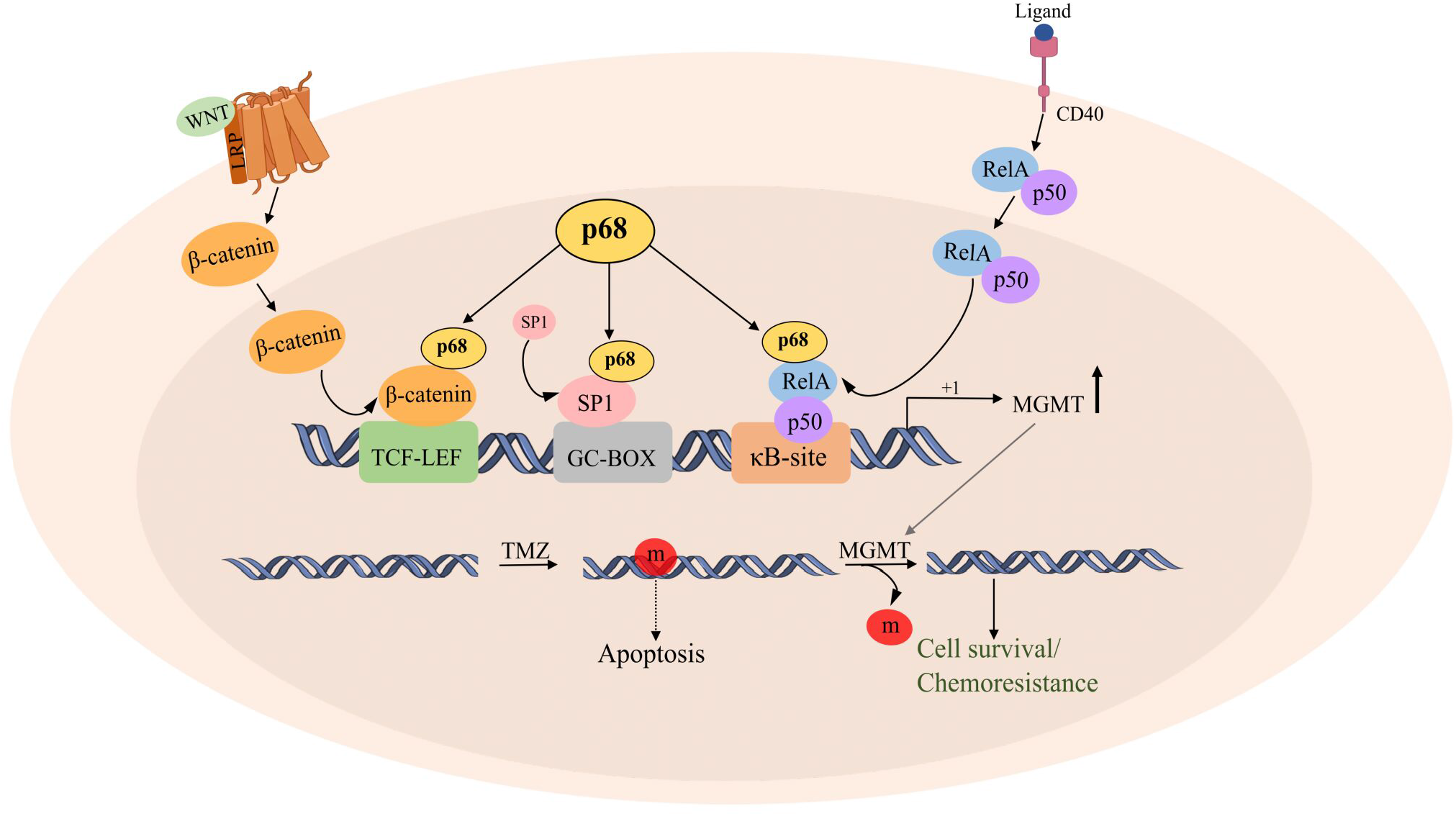
Proposed model for molecular mechanism of MGMT gene regulation. Molecular mechanisms defining DDX5 (p68) as a master regulator of MGMT *via* β-catenin/RelA/SP1 axis and subsequently manipulating cell survival/apoptosis in the context of TMZ mediated chemoresistance in colon cancer cells.

In summary, our work thus establishes a compelling link in controlling chemoresistance where MGMT, a master tumor aggressor is regulated by another principal oncogene and pro-survival molecule p68. Active MGMT enzyme works as a DNA repair protein, antagonising the effect of DNA alkylating agent, TMZ and support chemoresistance that further signals the cells to activate important cell cycle regulators and markers of oncogenesis and recurrence. This further complicates TMZ-based chemotherapy and results in dose up-modulation compromising patient health and treatment. Exploration of this novel p68-MGMT signalling might expectantly provide a new orchestration in the clinical management of colon cancer. This work can also consider p68 as a therapeutic target in colon cancer thus aiding us to develop a new therapeutic network.

## Material and methods

### Human colon tissue samples

Formalin-fixed paraffin-embedded sections, derived from post-surgical human colon carcinoma (n = 20) and adjacent normal colon (n = 5) tissues collected from Indian patients were used in this study. The samples were collected as per all medical and ethical regulations, including patient consent, and with formal approval from the institutional ethical committees of both CSIR-IICB and Park Clinic (source of colon carcinoma and normal samples).

### Histological analysis and immunohistochemistry (IHC)

Histological and Immunohistochemical studies were conducted as described before.^25–27^ For image scoring purposes, an overall H-score^28^ ranging from 100 to 300 was achieved where the degree of staining (0–100%) was multiplied by the intensity pattern of staining (set at 1: negative or weak, 2: moderate and 3: strong). The slides were viewed and images were captured at ×20 by EVOS XL Cell Imaging System (Life Technologies-Thermo Fischer Scientific).

### Bioinformatic analyses

A GeneMANIA server was used to check the functional relationship among the desired genes. 5 genes (DDX5, MGMT, SP1, RELA, CTNNB1) were assigned on Cytoscape and with the query question that is are all these 5 proteins presented on a common platform or not, we created a protein-protein interaction (PPI) network in Cytoscape that displays the functional relationships of proteins with their physical interaction, co-expression profile, colocalization potential, and genetic interaction.^29, 30^ TCGA mRNA normalized RSEM modules were used by Gene Set Cancer Analysis (GSCA) tool. The samples of COAD in GSCALite provide mRNA differential expression with paired tumor and normal samples.^31^ The fold change is calculated as mean (Tumor) / mean (Normal) and also the FDR was used to adjust the p-value. Those genes with fold change (FC > 2) and significance level (FDR > 0.05) were retained for the figure production. Next for analysing the relative transcriptional expression of DDX5, MGMT, CTNNB1, RELA and SP1 between tumor vs normal individuals’ data in Colon Adenocarcinoma (COAD), we used Gene Expression Profiling Interactive Analysis (GEPIA) where p < 0.01 was considered statistically significant.^32^ MGMT and DDX5 (p68) gene expressions were analysed in multiple cancers including colorectal carcinoma using UALCAN (http://ualcan.path.uab.edu) database.^33^ The transcriptional expression of MGMT and DDX5 was further checked in individual cancer stages of colon carcinoma. *P* value *<* 0.05 was considered significant.

### Expression plasmids

P68 was subcloned from pSG5-Myc vector into pIRES-hrGFP-1a, pGZ21dx and pGEX-4T1. pBI-β-catenin was sub-cloned into pGZ21dx vector. pCMV4 p65 (RelA) was purchased from Addgene. Human SP1 (-1302 to + 36) was cloned into pCDNA3.1 H (+) vector. pSV2-MGMT was a generous gift to our lab from Dr. Bernd Kaina, Nuclear Research Center Karlsruhe, Germany. The p-954/+24 ML and p-3500/+24 ML variants of human MGMT promoter subcloned in the pGL2 basic vector were kind gifts to our lab by Dr. Sankar Mitra, Houston Methodist Academic Institue, Houston, Texas, USA.

### Short hairpin RNA (shRNA) mediated knockdown

shRNA against β-catenin was purchased from Addgene (pLKO.1 puro shβ-catenin; Addgene cat no.18803). shRNA targeting RelA and SP1 were cloned into pLKO.1-Puro vector individually according to Addgene’s protocol. The DNA constructs were further verified by restriction digestion and sequencing. The list of the primers used is given in Table S1.

### Cell culture, transfections and treatments

HEK 293, HCT 116, SW480 and HT29 cells were cultured in Dulbecco’s modified eagle medium (DMEM) supplemented with 10% heat-inactivated fetal bovine serum (GIBCO-Invitrogen), 2000 units/L penicillin (Invitrogen), 2mg/L streptomycin (Invitrogen) and gentamycin (Invitrogen). Cells were cultured at 37°C in a humid incubator with a set atmosphere of 5% CO2. Transfection of cells was carried out using Lipofectamine 3000 (Invitrogen) following manufacturer’s protocol. In case of drug treatment following DNA transfection, drug treatment was done at least 24 hr post-transfections. FH535, Bay11 and Mithramycin A (Sigma Aldrich), the antagonists of β-catenin, RelA and SP1 pathways respectively, were administered at a concentration of 15μM, 10μM and 24nM correspondingly for 24 hr, upon standardization of the doses. MGMT inhibitor O6-Benzyl Guanine was purchased from Sigma-Aldrich and was used at a concentration of 2.5μM for 24 hr unless mentioned otherwise. Temozolomide was also purchased from Sigma-Aldrich.

### Site-Directed Mutagenesis

Deletions of SP1 sites on human MGMT promoter were performed using Quick-Change XL Site-Directed Mutagenesis kit (Agilent Technologies, USA).^34^ The mutated promoters were further cloned into pGL3 basic vector. All the constructs were verified by sequencing. Sequences of the primers are given in Table S1.

### Western Blotting (WB)

Preparation of whole cell lysates and IB analyses were performed as described before.^35^ The following primary antibodies were used: p68 (Abcam), MGMT (Abcam); β-catenin (SCBT) and GFP (SCBT) (SantaCruz Biotechnology); RelA (CST), SP1 (CST), Cyclin D1 (CST), XIAP (CST) Cleaved PARP (CST) and PARP (CST) (Cell Signaling Technology); Flag (SA), His (SA), Actin (SA), HRP-tagged anti-rabbit and anti-mouse secondary antibodies (SA) (Sigma-Aldrich). Densitometry values of the immunoblots were computed using GelQuant.Net software.

### RNA preparation and quantitative real-time PCR

Total RNA was extracted by using Trizol reagent (Invitrogen) following the manufacturer‘s protocol. cDNA was prepared and Real-time PCR was performed by using FastSYBR Green Master Mix (Applied Biosystems) in Via7 Real-Time PCR Instrument (Applied Biosystems) as described before.^34, 36^ Sequences of the primers are given in Table S1.

### Immunoblot analysis and IP assay

Whole-cell extracts were prepared following standard protocols.^25, 35^ Immunoblotting was performed following standard procedures.^37^ Antibodies used have been discussed before.

### GST-fusion protein purification and GST pull-down assay

Escherichia coli Rosetta (DE3) cells transformed with either GST (vector control) or GST-p68 were cultured till mid-log phase, followed by induction with Isopropyl β-D-1 thiogalactopyranoside (0.8 mM) at 22 °C for 6 h. Protein purification was carried out following procedures as described earlier.^37, 38^

### Luciferase assay

Cells were transiently transfected with pGL2-MGMT promoter luciferase reporter construct along with Renilla luciferase plasmid (pRL-TK) and respective gene constructs as per experimental interest and indicated in the relevant figure(s). The efficiency of transfection was normalized with Renilla luciferase expression. Luciferase activity was determined luminometrically in the Varioskan Flash Multimode Reader (Thermo Fisher Scientific, Waltham, MA, USA) by the dual luciferase assay system (Promega, Madison, WI,USA) following manufacturer’s instructions and as described before.^39^

### Chromatin Immunoprecipitation (ChIP) Assay

ChIP assay was conducted as described previously.^34, 40^ Sequences of the primers are given in Table S1.

### Cell Viability and Wound Healing assays

Cell viability assay using MTT and wound healing (scratch) assay on 70-80% confluent monolayer cells were conducted as described before.^35^

### Immunofluorescence microscopy

The standard protocol of immuno-staining was observed as described before^34^ using HCT 116 cells followed by staining with primary antibodies, p68 (Abcam), MGMT (Abcam), β-catenin (Santa Cruz), RelA (CST) and SP1 (CST) and fluorochrome-conjugated secondary antibodies (Alexa-Fluor). Hoechst was used as a nuclear counter stain. All slides were viewed and images were captured at ×20 using FluoView FV10i confocal laser scanning microscope (Olympus Life Science).

### Flow cytometric analysis

HCT 116 cells were transfected with 4μg of target vector or control vector, or treated with O6-BG or DMSO and then TMZ for 24 or 48 hrs as mentioned and harvested using Trypsin. The cells were further processed either by staining with Annexin V/Propidium Iodide (PI) for apoptotic assay or for cell-cycle analysis as described before^25, 35, 41^ and analysed in BD LSR-Fortessa using Flow-Jo Software (BD Biosciences).

### Colony formation assay

Treatments and transfections were performed on HCT 116 cells seeded at a density of 1 × 10^3^, as per the experimental design and with proper controls. After the respective time periods for transfections, cells were allowed to grow for 15 days in complete medium. The assay was performed as described previously.^42, 43^

### Statistical analyses

Paired Student‘s t-test was employed to determine the significance value in all experiments. The significance is presented as *P ≤ 0.05, **P ≤ 0.01, ***P ≤ 0.001 and ****P ≤ 0.0001, and nonsignificant differences are presented as NS. The differences in H-score values of all the concerned proteins between colon carcinoma and adjacent normal tissues were analysed by Mann–Whitney U test. All statistical analyses were performed using either SPSS (IBM) or GraphPad QuickCals software packages.

## Abbreviations

MGMT: O^6^-MethylGuanine-Methyl Transferase
SP1: Specificity Protein 1
O^6^-BG: O^6^-Benzyl Guanine
ChIP: Chromatin Immunoprecipitation
shRNA: Short Hairpin RNA
TMZ: Temozolomide
IB: Immunoblotting
Co-IP: Co-Immunoprecipitation
AFM: Atomic Force Microscopy.

## Acknowledgement

We are sincerely thankful to Dr. Bernd Kaina; Department of Genetics and Toxicology, Nuclear Research Center Karlsruhe, Germany, for kindly gifting us with the pSV2-MGMT construct, and to Dr. Sankar Mitra, Houston Methodist Academic Institue, Houston, Texas, USA, for generously gifting us the p-954/+24 ML and p-3500/+24 ML variants of human MGMT promoter subcloned in the pGL2 basic vector.

## Funding

This work is jointly supported by the Department of Science and Technology {NanoMission: DST/NM/NT/2018/105(G); SERB: EMR/2017/000992} and Focused Basic Research (FBR): MLP-142 and HCP-40, CSIR, Govt. of India to Dr. Mrinal K Ghosh.

## Conflicts of interest

The authors declare no conflicts of interest.

## Supplementary data

Supplementary figures S1 – S5; Table S1 – list of all primers used.

## Notes

### Competing Interest Statement

The authors have declared no competing interest.

