## Supplementary Figures S1-S5 for "DDX5 (p68) orchestrates β-catenin, RelA and SP1 mediated MGMT gene expression: Implication in TMZ chemoresistance"

### Supplementary figures and figure legends

**Fig. S1**

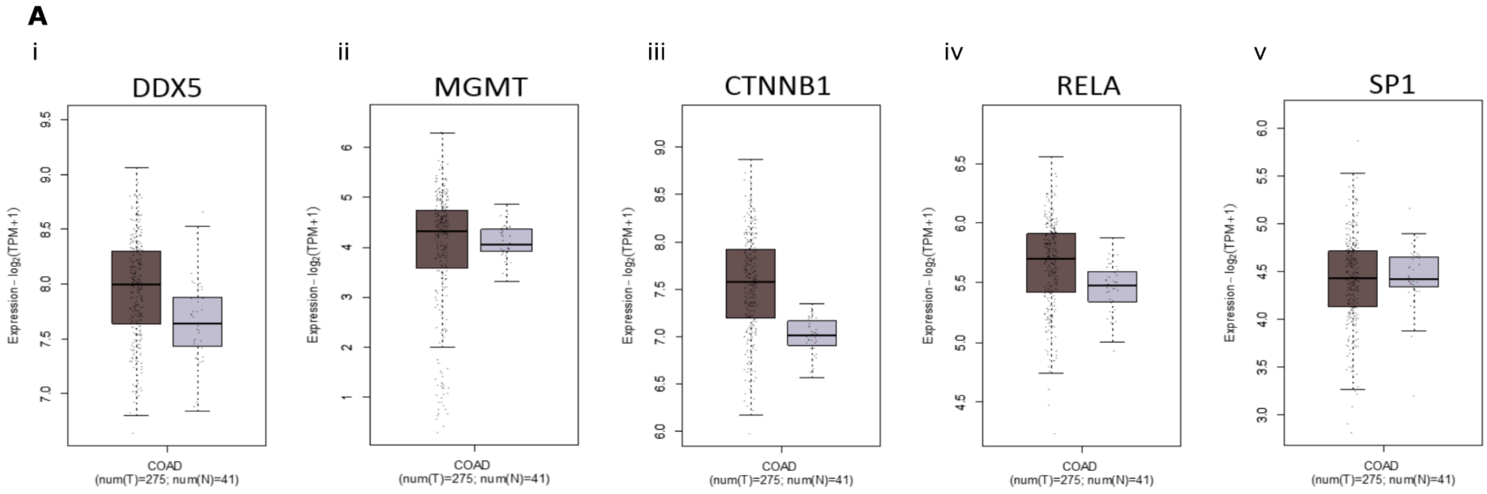

**Fig. S2**

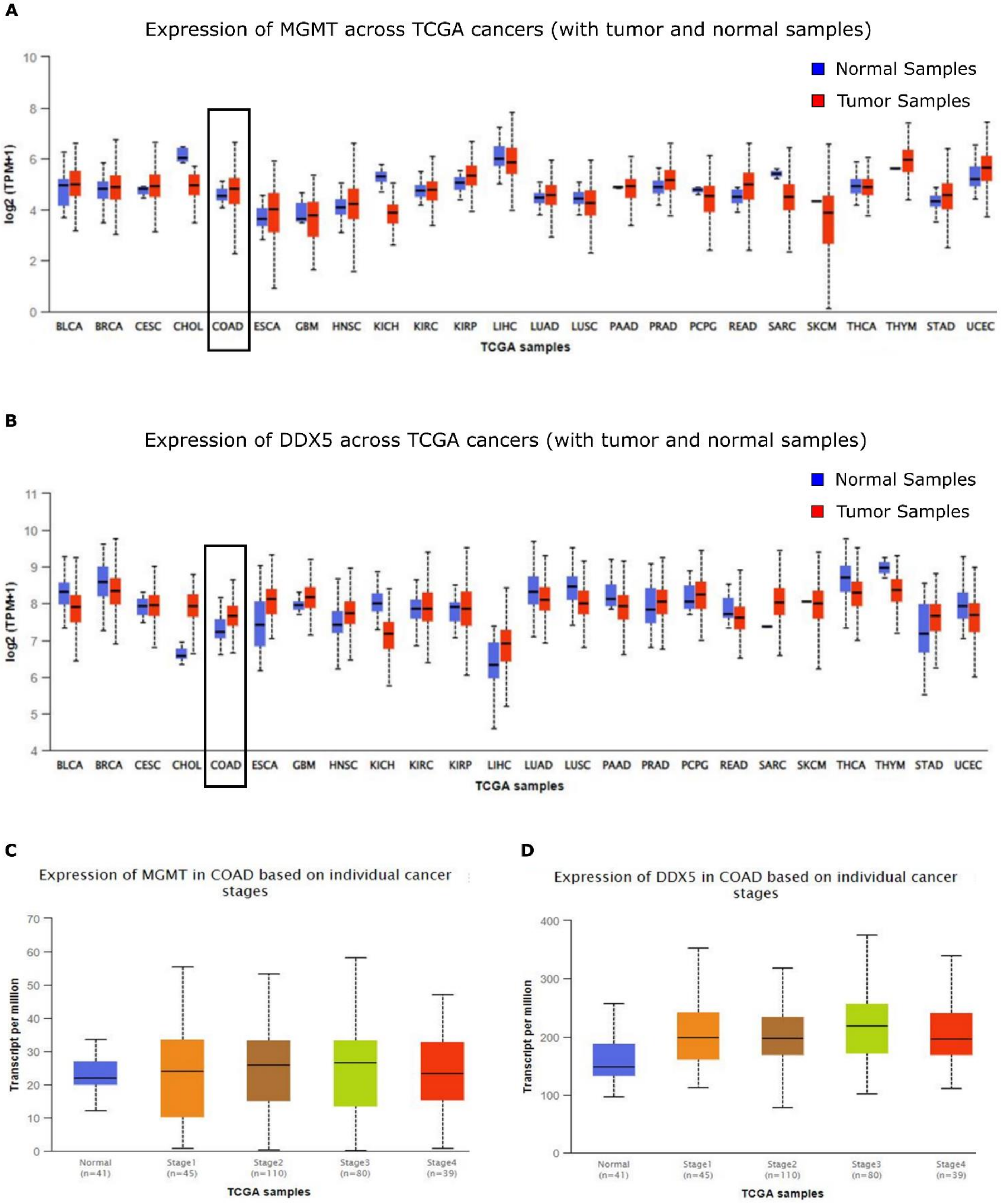
A
Expression of MGMT across TCGA cancers (with tumor and normal samples)
Normal Samples
Tumor Samples
log2 (TPM+1)
BLCA BRCA CESC CHOL COAD ESCA GBM HNSC KICH KIRC KIRP LIHC LUAD LUSC PAAD PRAD PCPG READ SARC SKCM THCA THYM STAD UCEC
TCGA samples
B
Expression of DDX5 across TCGA cancers (with tumor and normal samples)
Normal Samples
Tumor Samples
log2 (TPM+1)
BLCA BRCA CESC CHOL COAD ESCA GBM HNSC KICH KIRC KIRP LIHC LUAD LUSC PAAD PRAD PCPG READ SARC SKCM THCA THYM STAD UCEC
TCGA samples
C
Expression of MGMT in COAD based on individual cancer stages
Transcript per million
Normal (n=41)
Stage1 (n=45)
Stage2 (n=110)
Stage3 (n=80)
Stage4 (n=39)
TCGA samples
D
Expression of DDX5 in COAD based on individual cancer stages
Transcript per million
Normal (n=41)
Stage1 (n=45)
Stage2 (n=110)
Stage3 (n=80)
Stage4 (n=39)
TCGA samples

**Fig. S3**

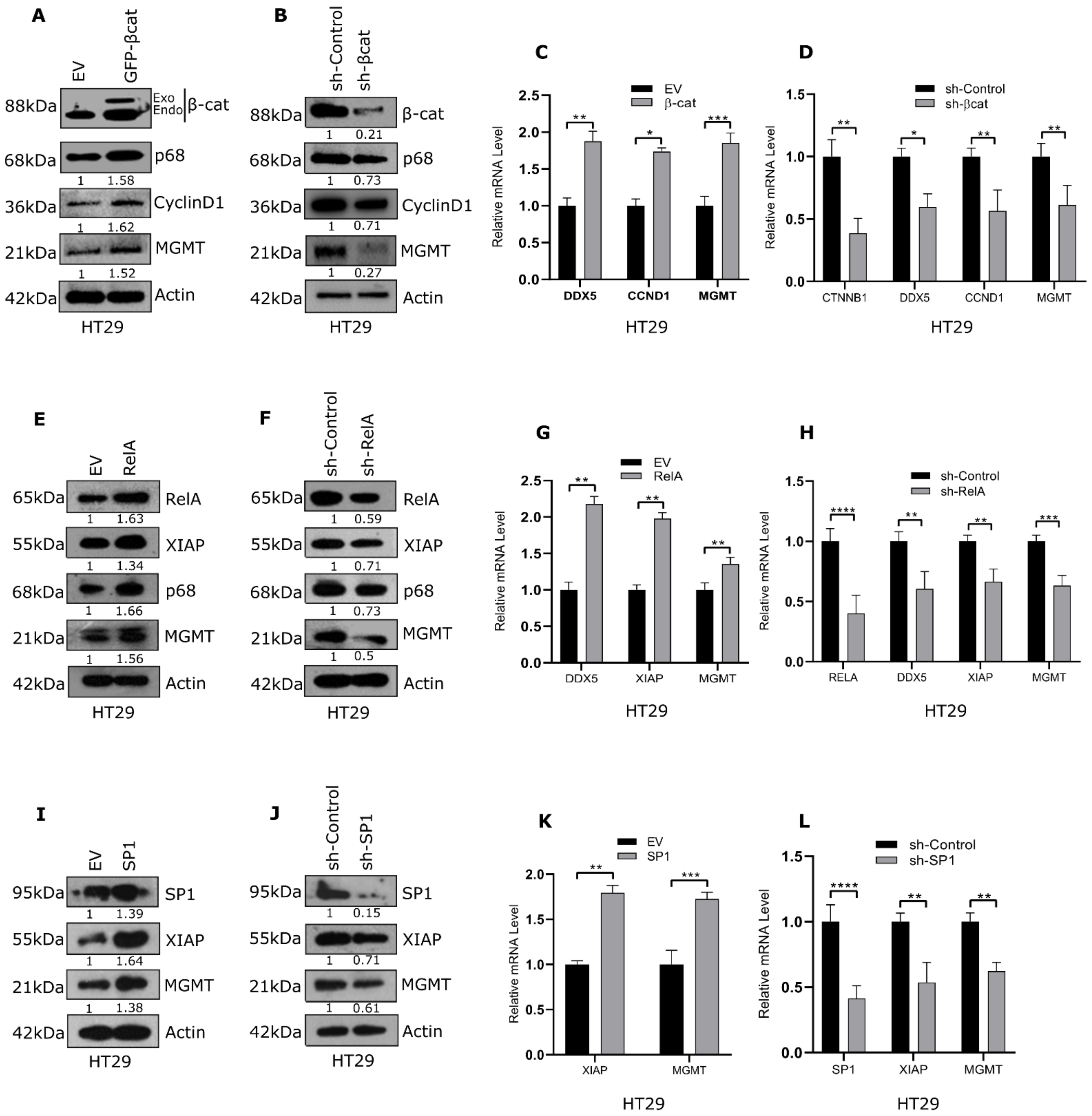

A
EV
GFP-βcat
88kDa
Exo
Endo
β-cat
68kDa
p68
1 1.58
36kDa
CyclinD1
1 1.62
21kDa
MGMT
1 1.52
42kDa
Actin
HT29
B
sh-Control
sh-βcat
88kDa
β-cat
1 0.21
68kDa
p68
1 0.73
36kDa
CyclinD1
1 0.71
21kDa
MGMT
1 0.27
42kDa
Actin
HT29
C
EV
β-cat
Relative mRNA Level
DDX5
CCND1
MGMT
HT29
D
sh-Control
sh-βcat
Relative mRNA Level
CTNNB1
DDX5
CCND1
MGMT
HT29
E
EV
RelA
65kDa
RelA
1 1.63
55kDa
XIAP
1 1.34
68kDa
p68
1 1.66
21kDa
MGMT
1 1.56
42kDa
Actin
HT29
F
sh-Control
sh-RelA
65kDa
RelA
1 0.59
55kDa
XIAP
1 0.71
68kDa
p68
1 0.73
21kDa
MGMT
1 0.5
42kDa
Actin
HT29
G
EV
RelA
Relative mRNA Level
DDX5
XIAP
MGMT
HT29
H
sh-Control
sh-RelA
Relative mRNA Level
RELA
DDX5
XIAP
MGMT
HT29
I
EV
SP1
95kDa
SP1
1 1.39
55kDa
XIAP
1 1.64
21kDa
MGMT
1 1.38
42kDa
Actin
HT29
J
sh-Control
sh-SP1
95kDa
SP1
1 0.15
55kDa
XIAP
1 0.71
21kDa
MGMT
1 0.61
42kDa
Actin
HT29
K
EV
SP1
Relative mRNA Level
XIAP
MGMT
HT29
L
sh-Control
sh-SP1
Relative mRNA Level
SP1
XIAP
MGMT
HT29

**Fig. S4**

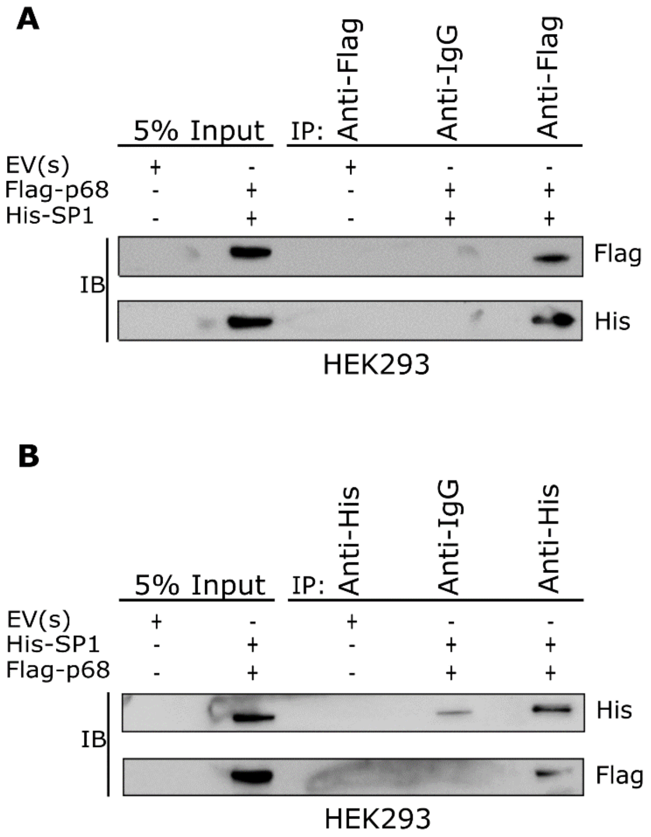
A
5% Input
IP:
Anti-Flag
Anti-IgG
Anti-Flag
EV(s) + - + - -
Flag-p68 - + - + +
His-SP1 - + - + +
IB
Flag
His
HEK293
B
5% Input
IP:
Anti-His
Anti-IgG
Anti-His
EV(s) + - + - -
His-SP1 - + - + +
Flag-p68 - + - + +
IB
His
Flag
HEK293

**Fig. S5**

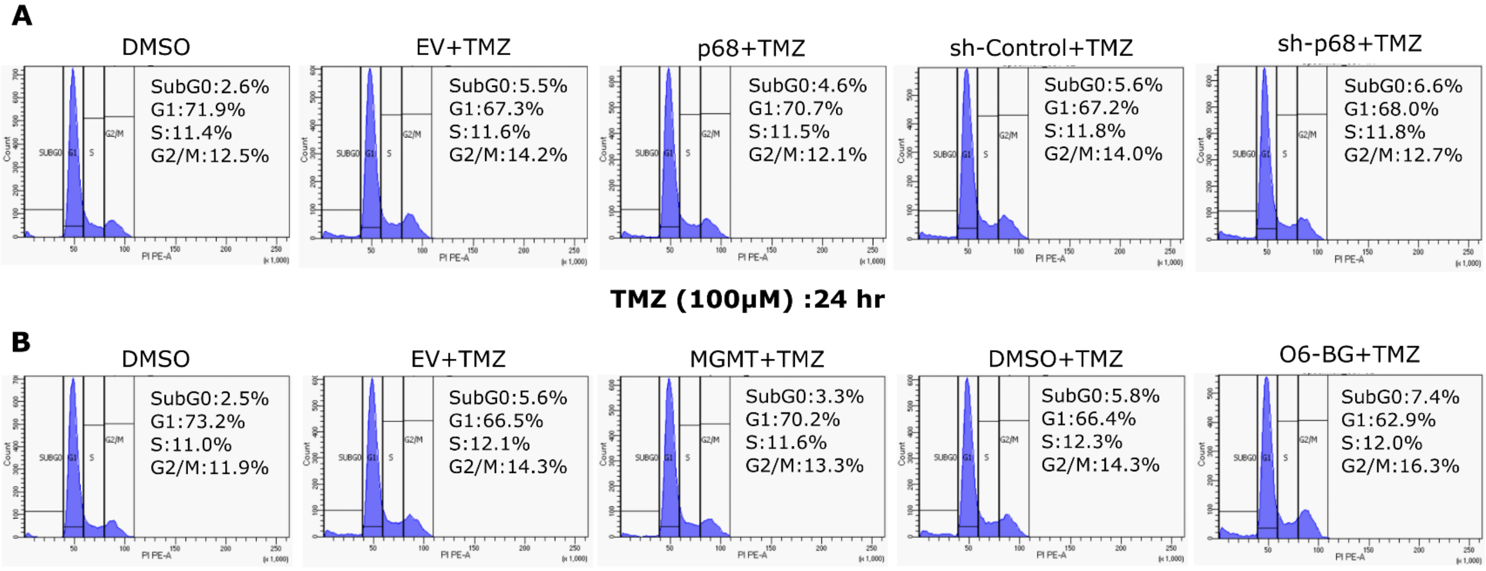
A
DMSO
SubG0:2.6%
G1:71.9%
S:11.4%
G2/M:12.5%
EV+TMZ
SubG0:5.5%
G1:67.3%
S:11.6%
G2/M:14.2%
p68+TMZ
SubG0:4.6%
G1:70.7%
S:11.5%
G2/M:12.1%
sh-Control+TMZ
SubG0:5.6%
G1:67.2%
S:11.8%
G2/M:14.0%
sh-p68+TMZ
SubG0:6.6%
G1:68.0%
S:11.8%
G2/M:12.7%
TMZ (100μM) :24 hr
B
DMSO
SubG0:2.5%
G1:73.2%
S:11.0%
G2/M:11.9%
EV+TMZ
SubG0:5.6%
G1:66.5%
S:12.1%
G2/M:14.3%
MGMT+TMZ
SubG0:3.3%
G1:70.2%
S:11.6%
G2/M:13.3%
DMSO+TMZ
SubG0:5.8%
G1:66.4%
S:12.3%
G2/M:14.3%
O6-BG+TMZ
SubG0:7.4%
G1:62.9%
S:12.0%
G2/M:16.3%

### Supplementary figure legends

**Fig. S1 Bioinformatics based transcriptional analysis of β-catenin, RelA, SP1, DDX5 and MGMT in Normal vs. Colon Cancer patients. (A)** GEPIA (Http://gepia.cancer-pku.cn) was used to compare the differential expression profile of 5 selected genes in COAD vs. Normal tissue samples.

**Fig. S2 Bioinformatic analysis of MGMT and DDX5.** Bioinformatics based Pan-Cancer analysis of **(A)** MGMT and **(B)** DDX5 expression across the cancers using the UALCAN (http://ualcan.path.uab.edu/analysis.html) platform. BLCA, bladder urothelial carcinoma; BRCA, breast carcinoma; CESC, cervical squamous cell carcinoma and endocervical adenocarcinoma; CHOL, cholangiocarcinoma; COAD, colon adenocarcinoma; ESCA, esophageal carcinoma; GBM, glioblastoma multiforme; HNSC, head and neck squamous cell carcinoma; KICH, Kidney chromophobe; KIRC, kidney renal clear cell carcinoma; KIRP, kidney renal papillary cell carcinoma; LIHC, liver hepatocellular carcinoma; LUAD, lung adenocarcinoma; LUSC, lung squamous cell carcinoma; PAAD, pancreatic adenocarcinoma; PRAD, prostate adenocarcinoma; PCPG, pheochromocytoma and paraganglioma; READ, rectum adenocarcinoma; SARC, sarcoma; SKCM, skin cutaneous melanoma; THCA, thyroid carcinoma; THYM, thymoma; STAD, stomach adenocarcinoma; UCEC, uterine corpus endometrial carcinoma. Relative mRNA expression was analysed for **(C)** MGMT and **(D)** DDX5 in different stages of COAD.

**Fig. S3 β-catenin, RelA and SP1 positively regulate protein and mRNA expression of MGMT gene in HT29 colon cancer cells.** Immunoblot analysis of cells transfected with either **(A)** GFP-β-catenin or the empty vector, **(E)** pCMV4-RelA or the empty vector and **(I)** pcDNA3.1-SP1 or the empty vector and harvested at a total of 48 hr post transfection. Immunoblot analysis of cells transfected with either control shRNA or shRNA against **(B)** β-catenin **(F)** RelA and **(J)** Sp1 and harvested 48 hr post transfection. Relative mRNA expression (w.r.t 18s rRNA) of MGMT assessed. qRT-PCR analysis of cells transfected with **(C)** GFP-β-catenin, **(G)** pCMV4-RelA and **(K)** pcDNA3.1-SP1. qRT-PCR analysis of cells transfected with either control shRNA or shRNA against **(D)** β-catenin, **(H)** RelA and **(L)** Sp1 and harvested 48 hr post transfection. Error bars represent the mean (±) S.D. of independent two-tailed Student's t-tests, where P<0.0001 is represented as **** for highly significant.

**Fig. S4 p68 co-localizes and physically interacts with SP1.** HEK293 cells were grown in 100 mm cell culture dishes and were co-transfected with 4 μg of pcDNA3.1-SP1 and 4 μg of pIRES-p68 or with the empty vectors. 48 hr post-transfection, whole cell lysates were prepared followed by IP with **(A)** anti-Flag antibody or **(B)** anti- His antibody and analysis of the indicated proteins by immunoblotting with protein specific antibodies. IP with normal rabbit immunoglobulin G (IgG) served as negative control. In all, 5% of the whole cell extracts used as input.

**Fig.S5 P68 actively regulates MGMT based TMZ response in HCT 116 cells. (A)** HCT 116 cells transfected with either pIRES-p68/control empty vector for p68 overexpression or with shRNA against p68/control shRNA for p68 knockdown and then 24 hr later treated with 100μM of TMZ for next 24 hr. Treated cells were processed and cell cycle analysis carried by flow cytometry. **(B)** HCT 116 cells transfected with pSV2-MGMT/control empty vector (PGZ) for MGMT overexpression or treatment with 25μM O6-BG/DMSO control for MGMT inhibition and then after 24 hr of transfection/treatment, treated with 100μM TMZ for 24 hr were then for cell cycle analysis by flow cytometry.

### Table S1. List of primers used in the study.

***Cloning***:

*RelA-shRNA:*

F-5'- CCGGGGGAAATACGTGGAGACACTACTCGAGTAGTGTCTCCACGTATTTCCCTTTTTG-3'

R-5'- AATTCAAAAAGGGAAATACGTGGAGACACTACTCGAGTAGTGTCTCCACGTATTTCCC-3'

*SP1*-shRNA:

F-5'-CCGGGCCAATAGCTACTCAACTACTCTCGAGAGTAGTTGAGTAGCTATTGGCTTTTTG
-3'
R-5'- AATTCAAAAAGCCAATAGCTACTCAACTACTCTCGAGAGTAGTTGAGTAGCTATTGGC
-3'

***qRT-PCR*:**

*DDX5:*

F-5’-TGAGCGACCTTATCTCTGTGC-3’

R-5’-CCTGGAACGACCTGAACCTC-3’

*MGMT:*

F-5’-ACCGTTTGCGACTTGGTACT-3’

R-5’-TGCTCACAACCAGACAGCTC-3’

*β-catenin:*

F-5’-TACCTCCCAAGTCCTGTATGAG-3’

R-5’-TGAGCAGCATCAAACTGTGTAG-3’

*Cyclin D1:*

F-5’-CCGTCCATGCGGAAGATC-3’

R-5’-GAAGACCTCCTCCTCGCACT-3’

*RelA*:

F-5’-ACAACAACCCCTTCCAAGTTCC-3’

R-5’-ACTGTCACCTGGAAGCAGAGC-3’

*SP1*:

F-5’-GTGGTGGTGCCTTTTCACAG-3’

R-5’-CAGCAGAGCCAAAGGGGATG-3’

*XIAP*:

F-5’-GACAGTATGCAAGATGAGTCAAGTCA-3’

R-5’-GCAAAGCTTCTCCTCTTGCAG-3’

*18S rRNA:*

F-5’-GCTTAATTTGACTCAACACGGGC-3’

R-5’-AGCTATCAATCTGTCAATCCTGTC-3’

***SDM*:**

*SP1*-prom-Δ1:

F-5’-GGGCGCGACCGGGTCGGCGC-3’

R-5’-GCGCCGACCCGGTCGCGCCC-3’

*SP1*-prom-Δ2:

F-5’-GCCGGTACAAGCCGCTGAGCCCGG-3’

R-5’- CCGGGCTCAGCGGCTTGTACCGGC-3’

***PCR in the ChIP Assay***

*TCF-4 TBE B1-ChIP:*

F-5’-AGCCTGTATTGTCACCAGGG-3’

R-5’-CTAAGCCACCATGCTACGGG-3’

*TCF-4 TBE B2-ChIP:*

F-5’-GGTCGGGCGGGAACA-3’

R-5’-TCACCAAGTCGCAAACGGT-3’

*RelA R1-ChIP:*

F-5’-CCGGTTCTAACTGGGTCCTG-3’

R-5’-CCCGGTGAAGTTTCCCTGTT-3’

*SP1 S1-ChIP:*

F-5’-TGACAGGAAAAGGTACGGGC-3’

R-5’- CAGACACTCACCAAGTCGCA -3’

*SP1 S2-ChIP:*

F-5’-GTCTAGGCCATCGGTGACTG-3’

R-5’-CCCGCTTAGTGAGAATCCCG-3’

*Cyclin D1-ChIP:*

F-5’-GTAACGTCACACGGACTACAGG-3’

R-5’-GCACACATTTGAAGTAGGACACC-3’

*XIAP-ChIP (RelA):*

F-5’-TGTTTCCGGTCCATCTGCTT-3’

R-5’-CCCCAGCTCCAGTGTTTCTT-3’

*XIAP-ChIP (SP1):*

F-5’-ACCGTGGGAAAAACCATCCTT-3’

R-5’-TTCAACTTACCACTTGGGCCG-3’

*Actin-ChIP:*

F-5’- TGCACTGTGCGGCGAAGC-3’

R-5’- TCGAGCCATAAAAGGCAA-3’
